# Abundance of Whales in West and East Greenland in Summer 2015

**DOI:** 10.1101/391680

**Authors:** Rikke G. Hansen, Tenna K. Boye, Rasmus S. Larsen, Nynne H. Nielsen, Outi Tervo, Rasmus D. Nielsen, Marianne H. Rasmussen, Mikkel H.S. Sinding, Mads Peter Heide-Jørgensen

## Abstract

An aerial line transect survey of whales in West and East Greenland was conducted in August-September 2015. The survey covered the area between the coast of West Greenland and offshore (up to 100 km) to the shelf break. In East Greenland, the survey lines covered the area from the coast up to 50 km offshore crossing the shelf break. A total of 423 sightings of 12 cetacean species were obtained and abundance estimates were developed for common minke whale, from now on called minke whale, (*Balaenoptera acutorostrata)* (32 sightings), fin whale (*Balaenoptera physalus*) (129 sightings), humpback whale (*Megaptera novaeangliae)* (84 sightings), harbour porpoise (*Phocoena phocoena)* (55 sightings), long-finned pilot whale, from now on called pilot whale, (*Globicephala melas)* (42 sightings) and white-beaked dolphins (*Lagenorhynchus albirostri)* (50 sightings). The developed at-surface abundance estimates were corrected for both perception bias and availability bias if possible. Data on surface corrections for minke whales and harbour porpoises were collected from whales instrumented with satellite-linked time-depth-recorders. Options for estimation methods are presented and the preferred estimates are: Minke whales: 5,095 (95% CI: 2,171-11,961) in West Greenland and 2,762 (95% CI: 1,160-6,574) in East Greenland, fin whales: 2,215 (95% CI: 1,017-4,823) in West Greenland and 6,440 (95% CI: 3,901-10,632) in East Greenland, humpback whales: 993 (95% CI: 434-2,272) in West Greenland and 4,223 (95%CI: 1,845-9,666) in East Greenland, harbour porpoise: 83,321 (95% CI: 43,377-160,047) in West Greenland and 1,642 (95% CI: 319-8,464) in East Greenland, pilot whales: 9,190 (95% CI: 3,635-23,234) in West Greenland and 258 (95% CI: 50-1,354) in East Greenland, white-beaked dolphins 15,261 (95% CI: 7,048-33,046) in West Greenland and 11,889 (95% CI: 4,710-30,008) in East Greenland. The abundance of cetaceans in coastal areas of East Greenland has not been estimated before, but the limited historical information from the area indicate that the achieved abundance estimates were remarkably high. When comparing the abundance estimates from 2015 in West Greenland with a similar survey conducted in 2007 there is a clear trend towards lower densities in 2015 for the three baleen whale species and white-beaked dolphins. Harbour porpoises and pilot whales however, did not show a similar decline. The decline in baleen whale and white-beaked dolphin abundance is likely due to emigration to the East Greenland shelf areas where recent climate driven changes in pelagic productivity may have accelerated favourable conditions for these species.

## INTRODUCTION

Most cetacean species that occur in West Greenland are subject to various levels of subsistence hunting where frequent abundance estimates are required for assessing the effects of the exploitation. Furthermore, the waters around Greenland are located in a climate sensitive area and large-scale changes in the marine environment may eventually influence the presence and abundance of whales in coastal areas of Greenland.

Aerial surveys for large whales have been conducted at regular intervals in West Greenland since 1984. The first two surveys in 1984 and 1985 were conducted with the intention of obtaining uncorrected line transect estimates of the abundance of common minke whales, from now on called minke whales, (*Balaenoptera acutorostrata*) and fin whales (*Balaenoptera physalus*), however, too few sightings were obtained to generate estimates. After 1985 surveys were conducted as combined cue counting and line transect surveys. Based on surveys conducted in 1987 and 1988 a cue counting estimate of 3,266 (cv=0.31) minke whales for both years combined was obtained. A survey in 1989 obtained too few sightings for any meaningful abundance estimate, however a survey in 1993 resulted in a cue counting estimate of 8,371 (cv=0.43) minke whales (Larsen 1995). A cue-counting estimate of 10,792 (cv=0.59) minke whales corrected for whales missed by the observers (perception bias) was obtained based on a survey conducted in 2005 (Heide-Jørgensen *et al.* 2008) and a survey in 2007 resulted in a fully corrected estimate of 16,609 (cv=0.41) minke whales (Heide-Jørgensen *et al.* 2010a).

The eight aerial surveys conducted between 1984 and 2007 each provided between 9 and 44 primary sightings of minke whales. Most sightings were of single individuals and sightings were widely dispersed on the banks of West Greenland. Given the difficulties in visually detecting minke whales it is unlikely that future surveys will obtain significantly more detections.

Surveys for fin whales have been conducted regularly between 1982 and 2007 in West Greenland but only three surveys were useful for abundance estimation (Heide-Jørgensen *et al.* 2008, 2010b). In 1987/88 fin whale abundance was estimated at 1,100 (cv=0.35) from an aerial cue counting survey (IWC 1992). In 2005, the abundance was estimated at 3,234 fin whales (cv=0.44) from an aerial line transect survey with independent observers that allowed for correction of perception bias but not availability bias (Heide-Jørgensen *et al.* 2008). A ship-based survey also conducted in 2005 gave a smaller abundance estimate (1,980 fin whales, cv=0.38) than the aerial survey (Heide-Jørgensen et al. 2007). The estimated abundance of fin whales from the aerial survey in 2007 (4,468 fin whales, cv=0.68) was corrected for perception bias but not for whales that were submerged during the survey (availability bias; Heide-Jørgensen *et al.* 2010b).

An estimate of 1,218 (cv=0.56) humpback whales (*Megaptera novaeangliae*) from a survey in West Greenland in 2005 was uncorrected for availability and perception biases (Heide-Jørgensen et al. 2008), but a survey in 2007 provided a fully corrected estimate of 2,704 humpback whales (cv=0.34, Heide-Jørgensen and Laidre 2015) with an estimate of a 9% annual rate of increase based on a time series of abundance estimates from 1984 to 2007 (Heide-Jørgensen *et al.* 2012).

The abundance of small cetaceans during summer has only been estimated once before in West Greenland. Hansen and Heide-Jørgensen (2013) provided fully corrected estimates of 8,133 (cv=0.41) long-finned pilot whales, from now on called pilot whales, (*Globicephala melas*), 11,984 (cv=0.19) white-beaked dolphins (*Lagenorhynchus albirostris*) and 33,271 (cv=0.39) harbour porpoises (*Phocoena phocoena*). All three species are hunted in Greenland and updates of abundance estimates are needed.

This study presents results from an aerial line transect survey of small and large cetaceans in East and West Greenland conducted in August and September 2015. Options for converting the at-surface density of whales to fully corrected total estimates of abundance are explored and applied to earlier partially corrected surveys. This requires the application of correction factors that adjust for whales missed by the observers and for whales that are not available to be detected at the surface.

## METHODS

### Data Collection

#### Aerial survey technique

An aerial line transect survey of whales was conducted from 18-25 August in East Greenland and between 27 August and 30 September 2015 in West Greenland. The survey platform was a Twin Otter, with long-range fuel tanks and two pairs of independent observers all with access to bubble windows. Sightings, survey conditions and a log of the cruise track (recorded from an external GPS) were recorded on a Geospatial Digital Video Recording System (Remote GeoSystems, Inc., Colorado, USA) that also allowed for continuous video recording of the trackline. Declination angle (*θ*) to sightings was measured when animals were abeam with Suunto inclinometers and converted to perpendicular distance (*x*) using the equation from Buckland et al. (2001): *X*= *V*\*tan(90-*θ*) where *v* is the altitude of the airplane. Time-in-view was calculated as the difference in time between the first detection and the time the sighting passed abeam. Target altitude and speed was 213 m and 167 km hr^−1^, respectively. Survey conditions were recorded by the primary observers at the start of the transect lines and whenever a change in sea state, horizontal visibility or glare occurred.

The survey was designed to systematically cover the area between the coast of West Greenland and offshore (up to 100 km) to the shelf break (i.e. the 200 m depth contour) by placing transects evenly across strata. Transect lines (n=124) in West Greenland were evenly placed in an east-west direction except for south Greenland where they were placed in a north-south direction (Fig. 1). In East Greenland 90 transects were designed to systematically cover the area from the coast over the shelf break up to 50 km offshore. The surveyed area was divided into 11 strata in West Greenland and 10 strata in East Greenland. Additional smaller strata covered selected fjord areas in West Greenland, where lines were placed following a zig-zag design when possible.

**Fig. 1.**
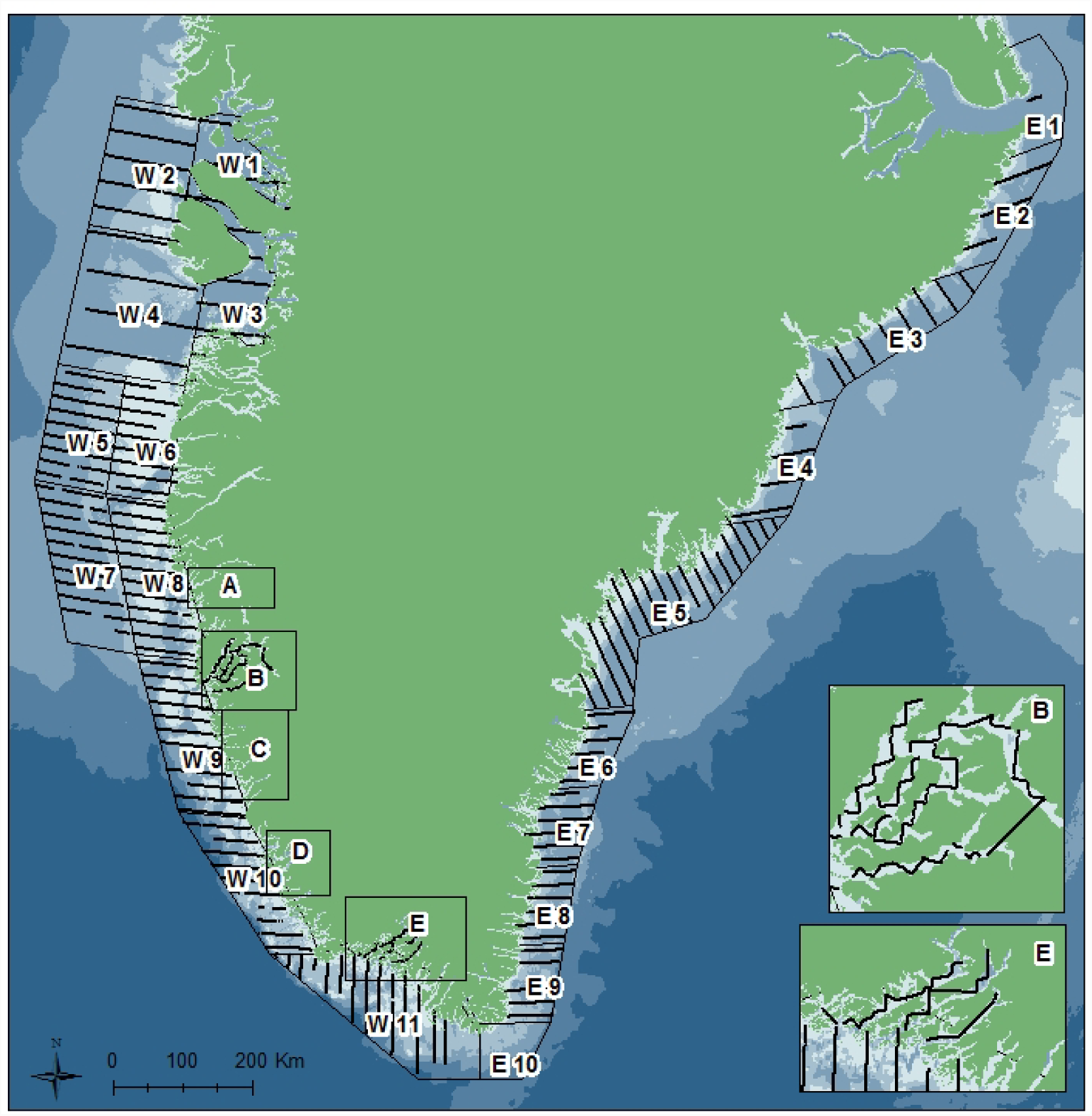
Survey effort in sea state <5 in East and West Greenland with delineation of strata and depth contours indicated (2,000m, 1,000 m, 200m and 100m). Block A (strata 12-18), B (strata 19), C (strata 20-25), D (strata 26-29) and E (strata 30-35). Block A, C and D were not surveyed.

#### Instrumentation of minke and fin whales and harbour porpoises with satellite tags

Five minke whales and two fin whales were tagged with satellite linked time-depth-recorders in July and August 2013-2017 (Table 1). The whales were pursued from an open skiff with outboard engine operating in the waters off the town of Maniitsoq (65°25’N and 52°54’W) in central West Greenland.

**Table 1.** Overview of data used for correction of the surface availability of the whales. Only data collected during daylight (07:00-19:00 GMT) and before 27 September are included. Data were extracted from messages relayed by satellite-linked time-depth-recorders.

| Species | Year | Month | n | 0 m | 0-2 m | 0-7m | Reference |
| --- | --- | --- | --- | --- | --- | --- | --- |
| **Minke whale**, West Greenland | - | - | - | - | - | - | - |
| #7934, | 2013 | 21/8-27/9 | 99 hrs | 2.21 | 16.5 | - | This study |
| #1328 | 2013 | 10/7-11/9 | 10 hrs | 1.38 | 23.3 | - | This study |
| #3964 | 2014 | 30/7-11/8 | 36 hrs | 0.16 | 12.15 | - | This study |
| #24642 | 2016 | 2/8 | 2 hrs | 0.25 | 12.4 | - | This study |
| #26715 | 2016 | 9-13/8 | 6 hrs | 8.55 | 21.3 | - | This study |
| Weighted mean (SE) | - | - | - | 1.90 (0.80) | 16.06 (1.30) | - | - |
| **Fin whale**, West Greenland |  |  |  |  |  |  |  |
| #26712 | 2016 | 11-22/8 | 15 hrs | 1.27 | 18.13 | - | This study |
| #168439 | 2017 | 23/7-8/9 | 239 hrs | 15.3 | 19.45 |  |  |
| Weighted mean (SE) | - | - | - | - | 19.37 (0.31) | - | - |
| **Humpback whale**, West Greenland | 2009-10 | May-July | 6 whales | - | 33.5 (3.4) | - | Heide-Jørgensen et al. 2015 |
| **Harbour porpoise**, West Greenland | - | - | - | - | - | - | - |
| #22854 | 2014 | Aug.- Sept. | ~720 hrs | 17.5 | - | - | This study |
| #93096 | 2014 | Aug.- Sept. | ~720 hrs | 26.5 | - | - | This study |
| #93100 | 2014 | Aug.- Sept. | ~720 hrs | 15.6 | - | - | This study |
| #93102 | 2014 | Aug.- Sept. | ~720 hrs | 20.0 | - | - | This study |
| #22849 | 2014 | Aug.- Sept. | ~720 hrs | 20.1 | - | - | This study |
| #22850 | 2014 | Aug.- Sept. | ~720 hrs | 18.9 | - | - | This study |
| #27261 | 2014 | Aug.- Sept. | ~720 hrs | 16.3 | - | - | This study |
| #27262 | 2014 | Aug.- Sept. | ~720 hrs | 17.7 | - | - | This study |
| #37235 | 2014 | Aug.- Sept. | ~720 hrs | 17.5 | - | - | This study |
| Mean (SE) | - | - | - | 19.0 (1.14) | - | - | - |
| **Pilot whale**, Faroe Isles | 2001 | Aug.- Sept. | 3 whales | - | - | 52.2 (0.002) | Heide-Jørgensen et al. 2002 |
| **Whitebeaked dolphin**, Iceland | 2006 | August | 14 hrs | - | 18 | - | Rasmussen et al. 2013 |

Instrumentation was conducted by using the Air Rocket Transmitter System (ARTS) initially developed by Heide-Jørgensen *et al.* (2001a, 2001b) and now widely used internationally for tagging baleen whales (Mate *et al.* 2007, Silva *et al.* 2013, Kennedy *et al.* 2014). The ARTS consists of an air gun with adjustable air pressure delivered by a scuba tank. The barrel of the ARTS is large enough to carry a plastic tube that acts as both a carrying rocket for the tag as well as a floatation device if the whale is missed. The ‘rocket’ is a cylinder with a finned tailpiece that provides stabilization during flight. The pressure and distance to the minke and fin whales when the rocket was launched was 12 bars and 5-10 m.

A cylindrical stainless steel implantable tag (22×113 mm, 1 AA cell, Mk10A Wildlife Computers, Redmond, WA) was used. It was equipped with triangular pointed steel arrow and had a transmission repetition period of 45s and a conductivity switch that prevented transmissions when the tag was underwater. The tags were not duty cycled, but were restricted to a maximum of 350 or 500 daily transmissions.

Nine live captured porpoises were instrumented with satellite-linked radio transmitters (SPLASH tags, Wildlife Computers) in 2014 with the same timing and locations as the minke whales (Nielsen et al. in press). The tag was attached to the dorsal fin using three 5 mm-diameter delrin nylon pins, that were pushed through holes drilled in the fin with a sterilized cork borer mounted on a cordless electric drill (Heide-Jørgensen et al. 2017, Nielsen et al. in press).

In order to collect data on the time spent at the surface, the satellite-linked dive recorders were equipped with a pressure transducer (WC ver. 10.2) and software (WC ver. 1.25p) that allowed for sampling of pressure every second. Information on the percentage of time spent at seven depth bins (TAD: 0, 0-1, 1-2, 2-3, 3-4, 4-5, >5m) were collected in 1hr intervals for the minke and fin whales. To correct for drift the pressure transducer were calibrated when breaking the surface by the salt-water switch that also controlled transmissions to the satellites. Information on the percentage of time spent between 0 and 2 m depth were collected in 1hr intervals for the harbour porpoises.

Averages of time spent at 0 and 0-2 m depths during daylight hours (8:00-18:00 local time) in August-September weighted by the sample sizes were calculated and used for developing availability correction factors (see below). The proportion of time spent at the surface for humpback whales was taken from Heide-Jørgensen *et al.* (2012), for pilot whales from Hansen and Heide-Jørgensen (2013) and for a white-beaked dolphin a tentative correction based on data from only one instrumentation was applied from Rasmussen *et al.* (2013).

### Data analysis

#### Mark-Recapture distance sampling correction for perception bias

The search method deployed used an independent observer configuration where the primary and secondary observers acted independently of each other. Detections of animals by the primary observer serve as a set of binary trials in which a success corresponds to a detection of the same group by the secondary observer on the same side. The converse is also true because the observers are acting independently; detections by secondary observers serve as trials for the primary observers. Analysis of the detection histories using logistic regression allows the probability that an animal on the trackline is detected by an observer to be estimated, and thus, abundance can be estimated without assuming *g*(0) is one, i.e. no perception bias. These methods combine aspects of both mark-recapture (MR) techniques and distance sampling (DS) techniques and so they are known as mark-recapture distance sampling (MRDS) methods (Laake and Borchers, 2004). Probability of detection is modelled as a function of the perpendicular distance from the track line including covariates like sea state, observer, side of plane and group size.

Although observers were acting independently, dependence of detection probabilities on unmodelled variables can induce correlation in the detection probabilities. Laake and Borchers (2004) and Borchers *et al.* (2006) developed estimators that assumed that detections were independent at zero perpendicular distance only – called point independence models - that are well suited for aerial surveys where no responsive movements are expected.

Sightings that were detected by both platforms (i.e. duplicates) were identified based on coincidence in timing (<3s), distance to sightings (± 200m), group size (± 20%) and species identification. In the few cases where different species were identified the more experienced front observer identifications were used. For duplicates identified with certainty a mean of group size and distance from both platforms were used for the density modelling.

Heterogeneity in the detection probabilities can be reduced by including explanatory variables in the MRDS model. The explanatory variables available in this survey were perpendicular distance to sighting, group size, sea state and observers and they were included in the MRDS models with model selection criteria to select the best model. Detection probability was estimated using the independent observer configuration implemented in Distance 6.2 (Thomas *et al.* 2010). Model selection and selection of either a uniform or half-normal detection function model was based on lowest AIC (Akaike Information Criteria).

Group abundance was estimated in each stratum using:

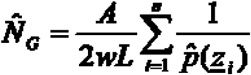

where *A* is the stratum area (in km^2^), *L* is the effort (in km) and *w* is the truncation distance (in m),_ is a vector of explanatory variables for group *i* (possibly including the group size,) and _ is the estimated probability of detecting group *i* obtained from the fitted MRDS model. Individual animal abundance is estimated by:

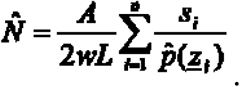

The estimated group size in the stratum is given by:

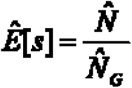

Effective search width was estimated at survey level (East and West Greenland combined) except for minke whales where strip width was estimated at global level (East and West Greenland separately). Encounter rates were estimated by stratum and detection probabilies were pooled across all strata. For each species it is estimated if group sizes were estimated at a global level (across all strata) in either East or West Greenland or at stratum level.

#### Perception bias in strip census estimates

The sample size for minke whales in West Greenland was too low to allow for distance sampling estimation and instead strip census estimation of density, with a constant probability of detection within a species-specific strip width, was used. The strip census estimate was developed with an average group size across all strata. A Chapman estimate was used to correct for perception bias ( ) by the observers:

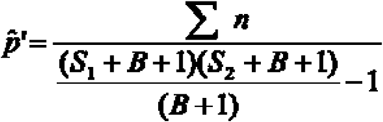

where *n* is the total number of sightings, *S*_*1*_ and *S*_*2*_ are the sightings by observer platform 1 and 2 only and *B* is the sightings by both platforms (Magnusson *et al.* 1978). Variance of ( ) was estimated with Jackknife methods.

Individual abundance in stratum *A* was developed from:

~~~
______
__________
~~~

where *G* is the average group size and *w* is the strip width.

#### Correction for non-instantaneous availability

Whales are available for detection for a short period of time during aerial surveys (i.e. some whales may be seen ahead of the plane). Therefore the probability that an animal was available to be seen was greater than the proportion of time it spends at depths at which it is visible. Laake et al. (1997) derived an equation for estimating the average probability of detecting a whale at the surface to correct for this:

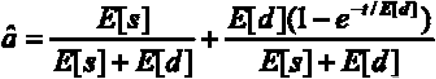

where *E*[*s*] is the average time the whale is at the surface, *E*[*d*] is the average time it is below the surface, and *t* is the window of time the whale is within visual range of the observers (‘time-in-view’).

Estimation of the availability correction factors and their variance was conducted by re-sampling the distributions of three parameters. Time-in-view (*t*) was resampled with replacement from the distribution of samples. Comparison of the distribution of time-in-view samples from East and West Greenland was done with a Kolmogorov-Smirnov test. Surfacing time (*st*) was sampled from a beta distribution with shape parameters and upper and lower limits that mimic a realistic distribution of the availability at the surface. The cue-rate per hour (*cr*) was assumed to have a normal distribution. *E*[*s*] and *E*[*d*] per hour were estimated as 3600 *st*/*cr* and 3600 (1-*st)*/*cr,* respectively.

It is assumed that the whales were only available for detection when they were close to the surface (0-2m) and that the proportion of time spent ( ) close to the surface was known from satellite linked-data recorders. In order to account for this availability bias, corrected abundance (denoted by the subscript ‘*c*’) was estimated by:

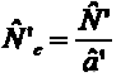

with estimated cv:

~~~
______
~~~

## RESULTS

*Estimation of time at surface and time in view for minke whale, fin whale and harbour porpoise*

The average time spent at all 7 sampled depth bins for the minke whales show that the largest proportion (percentage) of each hour was spent at depths >5 m and relatively small fractions of time were spent at the surface (Table 1).

The weighted average time spent between the surface and 2 m depth was around 16% for the five minke whales that provided dive data during day light hours (9:00-18:00) for July through 27 September in West Greenland.

The weighted average time spent between the surface and 2 m depth was around 19.4% for the two fin whales that provided dive data during day light hours (9:00-18:00) for the period 23 July through 8 September in West Greenland.

Nine harbour porpoises provided data on surfacing time (0 m) from August-September 2014 (Table 1). Data collected during daytime (07:00-19:00) were used and they were collected in 6 hr intervals providing around 720 hrs of measurements from each porpoise (2×6×60=720). The mean surfacing time of the nine whales was 19.0% (se=1.14).

### Estimation of abundance

A total of 423 sightings, covering 12 species of cetaceans and a few polar bear sightings, where obtained. Six species were detected in low numbers (n<12) leaving 6 species as candidates for abundance estimation.

*Minke whale abundance*

Sightings of minke whales were widely distributed in both East and West Greenland but there was a decline in numbers towards north with few or no sightings in the northern strata (Fig. 2). The highest density of minke whale sightings was found in the strata around the Kap Farvel (the southern tip of Greenland).

**Fig. 2.**
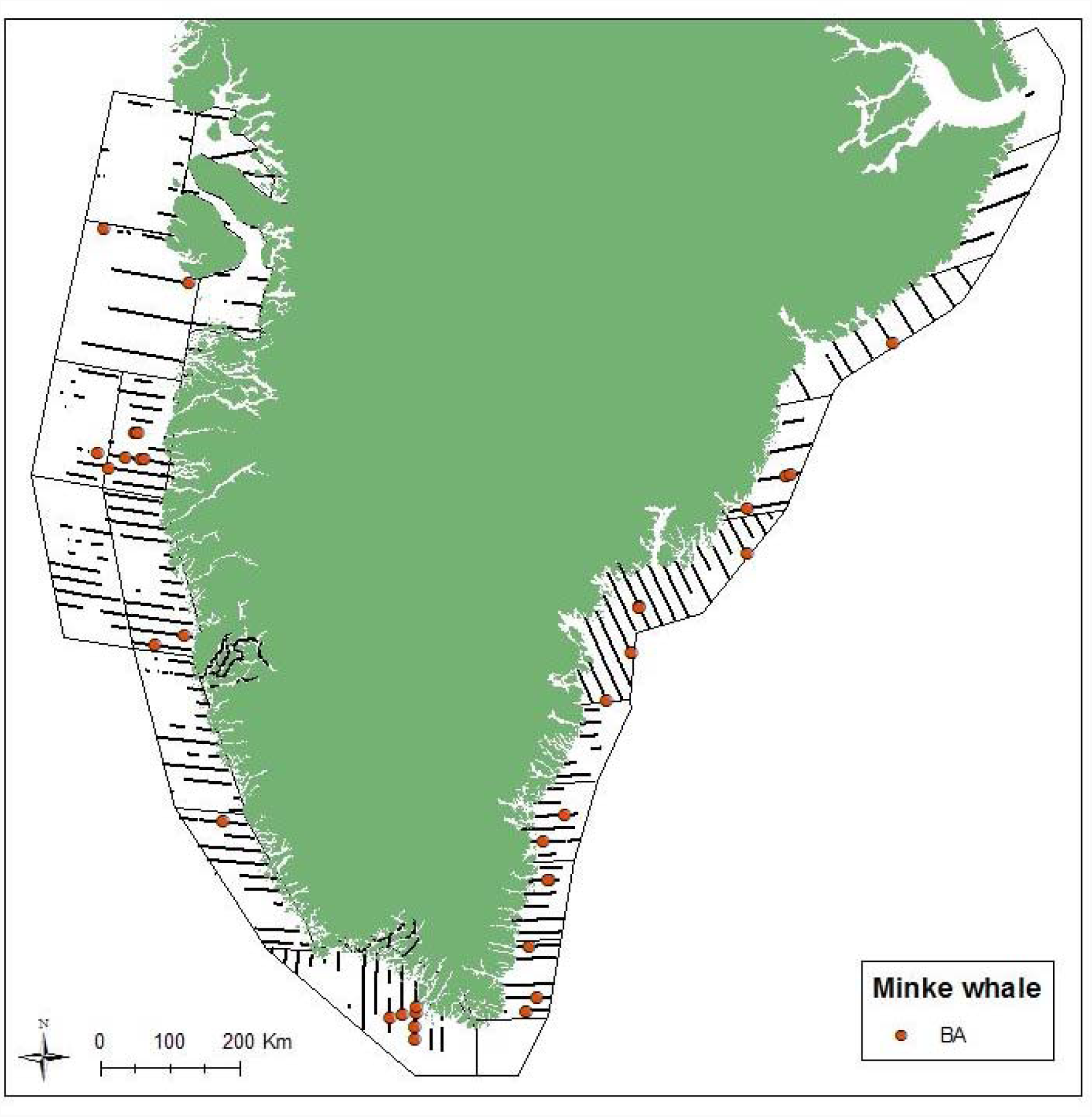
Sightings and survey effort in sea states <3 for minke whales in East and West Greenland.

The surveys in 2007 and 2015 in West Greenland had almost the same effort in both sea states <5 and <3 (Table 3), thus the number of sightings can be compared directly. The number of minke whale sightings in sea states<3 in 2015 was similar to the survey in 2007 and similar between the surveys in East and West Greenland despite twice the effort in West Greenland (Tables 2 and 3).

**Table 2.** Area of strata, effort and sightings.

| **2015 survey effort and sightings** | **East Greenland** | **West Greenland** |
| --- | --- | --- |
| **Sum of stratum area (km<sup>2</sup>)** | **114,742** | **220,924** |
| **Effort in ss<3 (km)** | **3,499** | **6,877** |
| Minke whale | 16 | 16 |
| Harbour porpoise | 2 | 53 |
| **Effort in ss<5 (km)** | **3,999** | **9,003** |
| Fin whale | 103 | 26 |
| Humpback whale | 61 | 23 |
| Pilot whale | 4 | 38 |
| White-beaked dolphin | 22 | 28 |
| Sperm whale | 3 | 5 |
| Bottlenose whale | 0 | 11 |
| Blue whale | 1 | 0 |
| Killer whale | 4 | 1 |
| Sei whale | 0 | 1 |
| Narwhal | 1 | 0 |
| Polar bear | 3 | 1 |

**Table 3.** Effort and number of sightings of humpback whales, fin whales, pilot whales, white-beaked dolphins, harbour porpoises and minke whales in West Greenland in 2007 and 2015.

| Strata | Area (km<sup>2</sup>) | Effort in km ss<5 |  | Sightings of humpback whale |  | Sightings of fin whale |  | Sightings of pilot whale |  | Sightings of white-beaked dolphin |  | Effort in km ss<3 |  | Sightings of harbour porpoise |  | Sightings of minke whale |  |
| --- | --- | --- | --- | --- | --- | --- | --- | --- | --- | --- | --- | --- | --- | --- | --- | --- | --- |
|  |  | 2007 | 2015 | 2007 | 2015 | 2007 | 2015 | 2007 | 2015 | 2007 | 2015 | 2007 | 2015 | 2007 | 2015 | 2007 | 2015 |
| W1 | 8,404 | 191 | 187 | 0 | 0 | 0 | 5 | 0 | 0 | 0 | 0 | 153 | 187 | 1 | 0 | 0 | 0 |
| W2 | 22,631 | 508 | 657 | 0 | 1 | 0 | 0 | 0 | 0 | 0 | 0 | 282 | 162 | 2 | 0 | 0 | 0 |
| W3 | 14,653 | 532 | 460 | 0 | 1 | 1 | 0 | 0 | 0 | 0 | 0 | 274 | 299 | 1 | 1 | 1 | 0 |
| W4 | 34,272 | 545 | 617 | 1 | 0 | 3 | 1 | 0 | 5 | 0 | 0 | 360 | 438 | 1 | 6 | 3 | 1 |
| W5 | 16,226 | 863 | 837 | 3 | 0 | 1 | 0 | 12 | 2 | 0 | 0 | 478 | 183 | 3 | 1 | 2 | 1 |
| W6 | 14,902 | 973 | 738 | 0 | 1 | 1 | 0 | 0 | 1 | 0 | 0 | 621 | 447 | 3 | 3 | 4 | 5 |
| W7 | 22,085 | 551 | 978 | 2 | 1 | 2 | 4 | 2 | 14 | 0 | 1 | 439 | 543 | 2 | 9 | 0 | 0 |
| W8 | 20,264 | 1,345 | 1,171 | 5 | 0 | 4 | 5 | 0 | 0 | 1 | 6 | 540 | 852 | 5 | 3 | 0 | 2 |
| W9 | 20,334 | 998 | 1,039 | 4 | 3 | 4 | 7 | 2 | 0 | 9 | 6 | 692 | 546 | 7 | 9 | 7 | 0 |
| W10 | 15,951 | 932 | 692 | 3 | 9 | 2 | 0 | 0 | 11 | 27 | 5 | 741 | 561 | 7 | 14 | 1 | 1 |
| W11 | 24,085 | 1,194 | 949 | 2 | 4 | 2 | 4 | 1 | 5 | 25 | 10 | 580 | 644 | 5 | 6 | 9 | 6 |
| W12 | 330 | 50 | 0 | 0 | 0 | 0 | 0 | 0 | 0 | 0 | 0 | 35 | 0 | 2 | 0 | 0 | 0 |
| W13 | 92 | 29 | 0 | 0 | 0 | 0 | 0 | 0 | 0 | 0 | 0 | 18 | 0 | 3 | 0 | 0 | 0 |
| W14 | 189 | 45 | 0 | 1 | 0 | 0 | 0 | 0 | 0 | 0 | 0 | 15 | 0 | 1 | 0 | 0 | 0 |
| W16 | 287 | 85 | 0 | 0 | 0 | 0 | 0 | 0 | 0 | 0 | 0 | 85 | 0 | 0 | 0 | 0 | 0 |
| W17 | 1,042 | 48 | 0 | 0 | 0 | 0 | 0 | 0 | 0 | 0 | 0 | 39 | 0 | 1 | 0 | 0 | 0 |
| W18 | 348 | 14 | 0 | 0 | 0 | 0 | 0 | 0 | 0 | 0 | 0 | 14 | 0 | 1 | 0 | 0 | 0 |
| W19 | 2,731 | 325 | 474 | 0 | 10 | 0 | 0 | 0 | 0 | 0 | 0 | 258 | 474 | 0 | 1 | 0 | 0 |
| W23 | 579 | 78 | 0 | 0 | 0 | 0 | 0 | 0 | 0 | 0 | 0 | 78 | 0 | 1 | 0 | 0 | 0 |
| W30 | 286 | 49 | 32 | 0 | 0 | 0 | 0 | 0 | 0 | 0 | 0 | 49 | 0 | 0 | 0 | 0 | 0 |
| W31 | 1,234 | 78 | 222 | 0 | 2 | 0 | 0 | 0 | 0 | 0 | 0 | 78 | 165 | 1 | 0 | 0 | 0 |
| W32 | 260 | 0 | 55 | 0 | 0 | 0 | 0 | 0 | 0 | 0 | 0 | 0 | 0 | 0 | 0 | 0 | 0 |
| **Total** | **221,185** | **9,433** | **9,108** | **21** | **23** | **24** | **26** | **17** | **38** | **62** | **28** | **5,829** | **5,501** | **47** | **53** | **27** | **16** |

The distribution of perpendicular distances of sightings (Figs 3 and 4) showed a large proportion of sightings close to the trackline indicating that there was not a blind spot for observers beneath the plane.

**Fig. 3.**
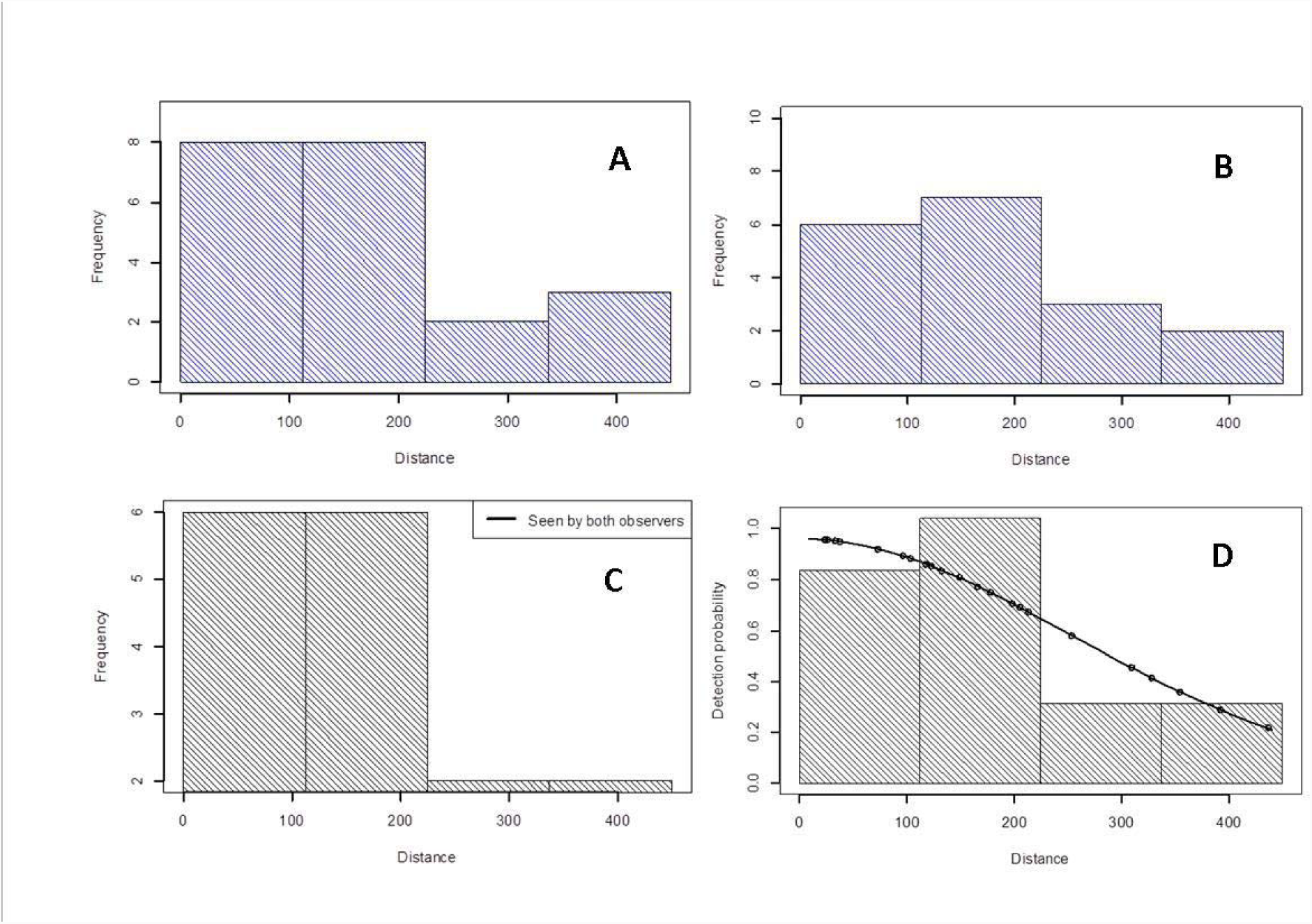
Scaled histograms of perpendicular distances (A, B, C) and the predicted detection probability functions (D) for minke whale groups. A) Front observers, B) rear observers, C) both observers and D) pooled detections with sightings fitted to a half-normal model where each observation prediction is shown as a circle. Distance is measured in meters from the line directly below the aircraft.

**Fig. 4.**
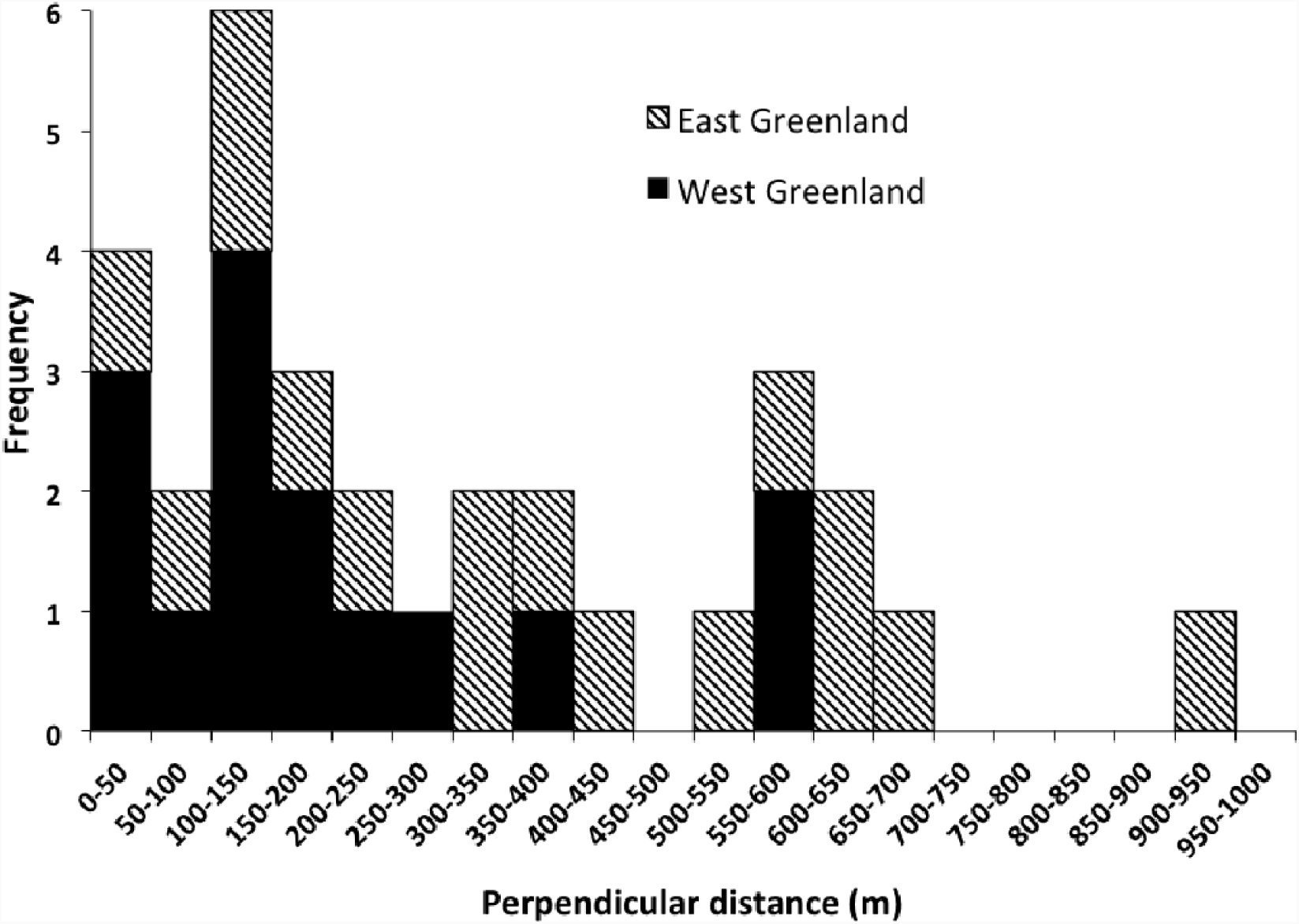
Distribution of sightings of minke whales in East and West Greenland in 2015 (n=31) in sea states <3.

All minke whale detections were of single individuals and the explanatory variables available to be included in the MRDS models were a) perpendicular distance to sightings, b) sea state and c) observer. Effort and sightings made in sea state above 2 was excluded from the model (Fig. 3). Estimates of correction factors for perception bias were developed for both MRDS methods and by the Chapman estimator for the strip census estimates (Table 4).

**Table 4.** Number of sightings seen by each observer and the number of duplicates (seen by both observers). The ‘Total’ column shows the number of sightings seen by observer 1 and observer 2 with the sightings seen by both removed. CV’s are given in parenthesis.

| Sea state | Observer 1 | Observer 2 | Seen by both | Total |
| --- | --- | --- | --- | --- |
| Minke whales East and West Greenland, dist<450 m (MRDS) |  |  |  |  |
| In ss<3 | 21 | 18 | 15 | 23 |
| Minke whales West Greenland, dist<300m |  |  |  |  |
| In ss<3 | 10 | 10 | 8 | 12 |
| Fin whales East and West Greenland (MRDS) |  |  |  |  |
| In ss<5 | 70 | 64 | 59 | 75 |
| Humpback whale East and West Greenland (MRDS) |  |  |  |  |
| In ss<5 | 72 | 57 | 53 | 76 |
| Harbour porpoise East and West Greenland (MRDS) |  |  |  |  |
| In ss<3 | 30 | 34 | 16 | 48 |
| Pilot whale East and West Greenland (MRDS) |  |  |  |  |
| In ss<5 | 30 | 29 | 27 | 32 |
| White-beaked dolphin East and West Greenland (MRDS) |  |  |  |  |
| In ss<5 | 42 | 42 | 36 | 48 |

Time-in-view for minke whales was on average 2.9 s for detection distances <450 m from the track line and 2.4 s for detections <300 m from the track line but the distribution were not significantly different (K-S, *p*=1), thus a common correction for availability bias (<450m) was applied (Table5).

In order to maintain consistency with the survey conducted in 2007 in West Greenland using identical methods (Heide-Jørgensen 2010a), separate analyses were chosen for East and West Greenland. The low number of sightings (n=12) precluded an MRDS analysis of minke whales in West Greenland alone but a strip census estimate (truncated at 300m), similar to that used in 2007, provided a fully corrected estimate for West Greenland of 5,095 (95% CI: 2,171-11,961) minke whales. About half the abundance of minke whales in West Greenland was found in the southernmost stratum next to the East Greenland survey area (Table 6).

**Table 6.**
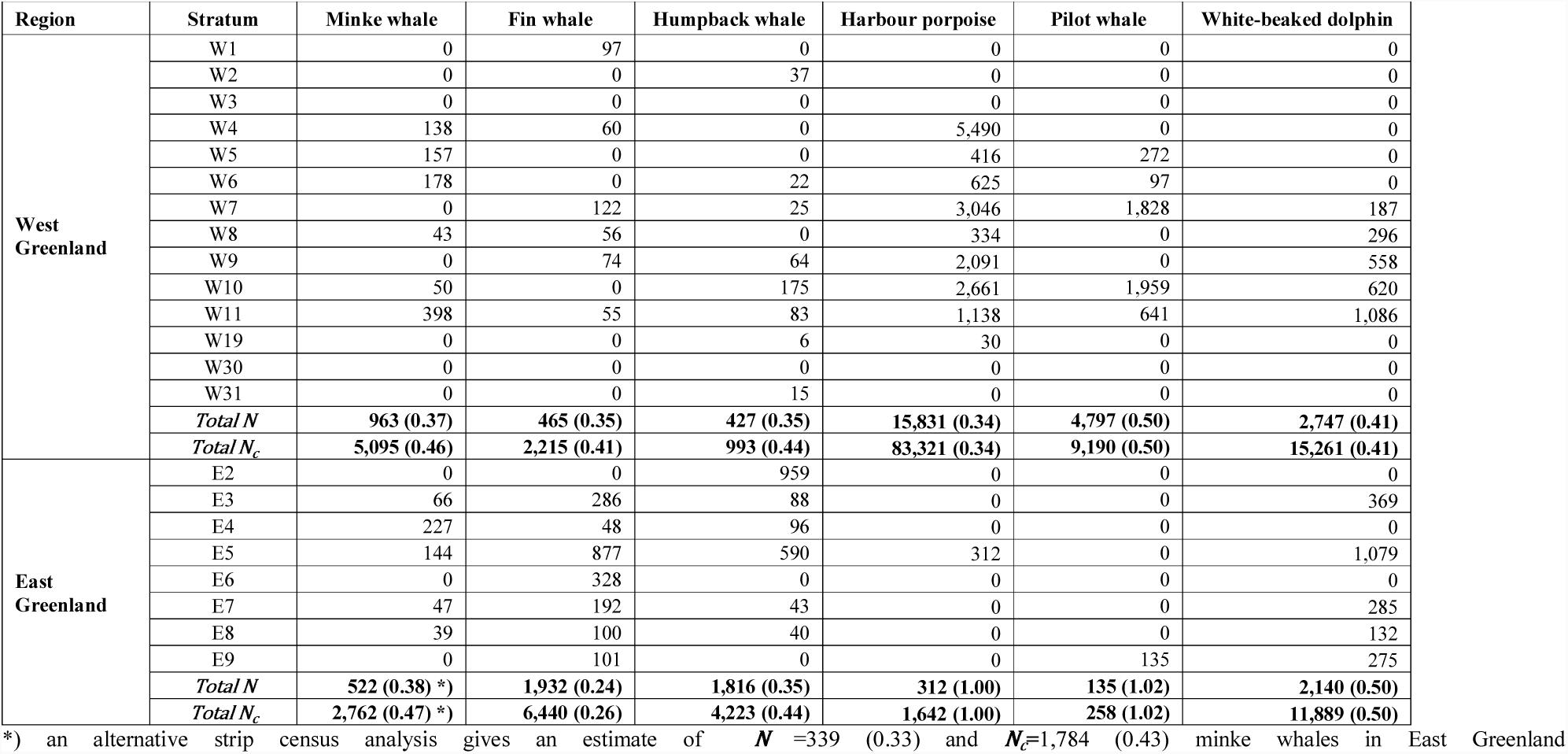
Abundance (*N*) per strata of minke whales at ss<3 (23 sightings), abundance of finwhales at ss<5 (75 sightings), humpback whales at ss<5 (76 sightings), harbour porpoise at ss<3 (76 sightings), pilot whales at ss<5 (42 sightings) and white-beaked dolphins at ss<5 (48 sightings) in 2015 by stratum in West and East Greenland estimated by MRDS analysis, except for minke whakes in West Greenland that are estimated by a strip census truncated at 300 m. Selected values are from selected estimates marked in bold text in Table 7. Availability correction factors from Table 5 were used to derive fully corrected abundance estimates (N_c_).

An MRDS analysis based on sightings combined from both East and West Greenland, truncated at 450 m and at sea state<3, excluded 4 observations, and provided a fully corrected (including availability bias) MRDS estimate of 2,762 (95% CI: 1,160-6,574) minke whales for East Greenland. An alternative analysis using only sightings from East Greenland and applying strip census estimation with an assumed strip width of 450m gives an estimate corrected for availability bias of 1,784 whales (cv= 0.43, 95% CI: 796-4,000). This estimate could not be reliably corrected for perception bias because all sightings were seen by observer 1 (hence there was no missing re-sightings for the front observer which seems unlikely based on previous double observer minke whale surveys).

A revised estimate of the abundance of minke whales in West Greenland in 2007 based on a previous aerial survey (Heide-Jørgensen et al. 2010a) using the availability factor applied to the survey in 2015 provided a new abundance estimate of 9,066 minke whales (95% CI: 4,333-18,973, Table 7).

**Table 7.** Fully corrected abundance estimates for baleen whales, harbour porpoise, pilot whales and white-beaked dolphins in West and East Greenland in 2005, 2007 and 2015. ESW is the effective search width and *N* is the number of sightings. All estimates are based on mark recapture distance methods except for the two minke whales estimates in West Greenland where strip census estimates with uniform detection functions were used due to the low number of sightings. Lcl and Ucl is lower and upper 95% confidence limits. E+W indicate that sightings from both East and West Greenland were used for deriving the detection functions.

| Method | ESW or strip width | Sightings | Model for perception bias | Perception bias | Detection depth | Availability bias incl. time-in-view | Estimate | cv | Lcl | Ucl |
| --- | --- | --- | --- | --- | --- | --- | --- | --- | --- | --- |
| Minke whale 2015 East Greenland | 450 m | 23 E+W | MRDS | 0.97 (0.04) | 0-2 m | 0.19 (0.27) | 2,762 | 0.47 | 1,160 | 6,574 |
| Minke whale 2015 East Greenland | 450 m | 10 | na | na | 0-2 m | 0.19 (0.27) | 1,784 | 0.43 | 796 | 4,000 |
| Minke whale 2015 West Greenland | 300 m | 12 | Chapman | 0.94 (0.06) | 0-2 m | 0.19 (0.27) | 5,095 | 0.46 | 2,171 | 11,961 |
| Minke whale 2007 West Greenland | 240 m | 18 | Chapman | 0.98 (0.02) | 0-2 m | 0.21 (0.25) | 9,066 | 0.39 | 4,333 | 18,973 |
| Fin whale 2015 East Greenland | 700 m | 75 E+W | MRDS | 0.99 (0.007) | 0-2 m | 0.30 (0.10) | 6,440 | 0.26 | 3,901 | 10,632 |
| Fin whale 2015 West Greenland | 700 m | 75 E+W | MRDS | 0.99 (0.007) | 0-2 m | 0.21 (0.22) | 2,215 | 0.41 | 1,017 | 4,823 |
| Fin whale 2007 West Greenland | 50-800 m | 24 | MRDS | 0.86 (0.09) | 0-2 m | 0.28 (0.22) | 15,957 | 0.72 | 4,531 | 56,202 |
| Fin whale 2005 West Greenland | 2500 m | 74 | MRDS | 0.51 (0.21) | 0-2 m | 0.33 (0.43) | 9,800 | 0.62 | 3,228 | 29,751 |
| Humpback whale 2015 East Greenland | 50-1200 | 76 E+W | MRDS global g | 0.98 (0.02) | 0-2 m | 0.43 (0.27) | 4,223 | 0.44 | 1,845 | 9,666 |
| Humpback whale 2015 West Greenland | 50-1200 | 76 E+W | MRDS global g | 0.98 (0.02) | 0-2 m | 0.43 (0.27) | 993 | 0.44 | 434 | 2,272 |
| Harbour porpoise 2015 East Greenland | 250 m | 48 E+W | MRDS | 0.81 (0.21) | 0 m | 0.19 (0.06) | 1,642 | 1.00 | 319 | 8,464 |
| Harbour porpoise 2015 West Greenland | 250 m | 48 E+W | MRDS | 0.81 (0.21) | 0 m | 0.19 (0.06) | 83,321 | 0.34 | 43,377 | 160,047 |
| Harbour porpoise 2007 West Greenland | 300 m | 31 | MRDS | 0.57 (0.23) | 0 m | 0.19 (0.06) | 54,284 | 0.36 | 27,627 | 106,664 |
| Pilot whales 2015 East Greenland | 700 m | 32 E+W | MRDS stratum g | 0.98 (0.03) | 0-7 m | 0.52 (0.002) | 258 | 1.02 | 50 | 1,354 |
| Pilot whales 2015 West Greenland | 700 m | 32 E+W | MRDS stratum g | 0.98 (0.03) | 0-7 m | 0.52 (0.002) | 9,190 | 0.50 | 3,635 | 23,234 |
| White-beaked dolphin 2015 East Greenland | 700 m | 48 E+W | MRDS | 0.94 (0.10) | 0-2 m | 0.18 (0.00) | 11,889 | 0.50 | 4,710 | 30,008 |
| White-beaked dolphin 2015 West Greenland | 700 m | 48 E+W | MRDS | 0.94 (0.10) | 0-2 m | 0.18 (0.00) | 15,261 | 0.41 | 7,048 | 33,046 |

#### Fin whale abundance

There were a few scattered observations of fin whales in West Greenland whereas large numbers were detected on the southern strata in East Greenland (Fig. 5).

**Fig. 5.**
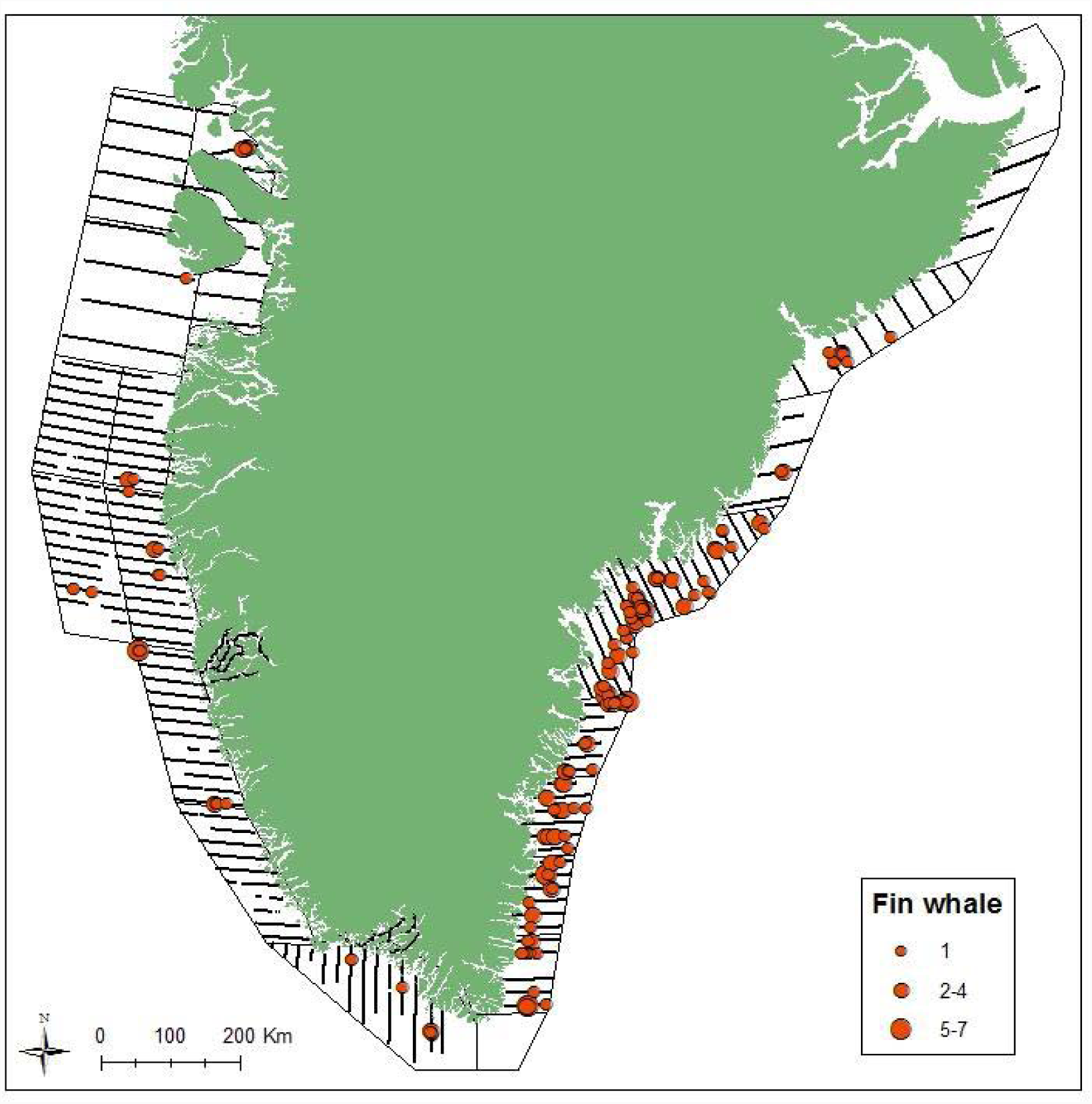
Survey effort in sea states <5 and sightings with group sizes of fin whales in East and West Greenland.

The number of sightings of fin whales in sea states <5 was about 4 times higher in East Greenland compared to West Greenland. The number of sightings in West Greenland was similar to the survey in 2007 (Tables 2 and 3).

The MRDS analysis was right truncated at 700 m leaving 75 observations for analysis with an overall expected group size of 1.2 (cv=0.09) fin whales in East and West Greenland combined. The explanatory variables available to be included in the MRDS models were a) perpendicular distance to sightings; b) group size, c) sea state and d) observer. Effort and sightings made in sea state above 4 were excluded from the model (Fig. 6). The best model, based on the lowest AIC, did not include any variables. There was little perception bias in the combined East and West Greenland data (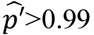, Table 4).

**Fig. 6.**
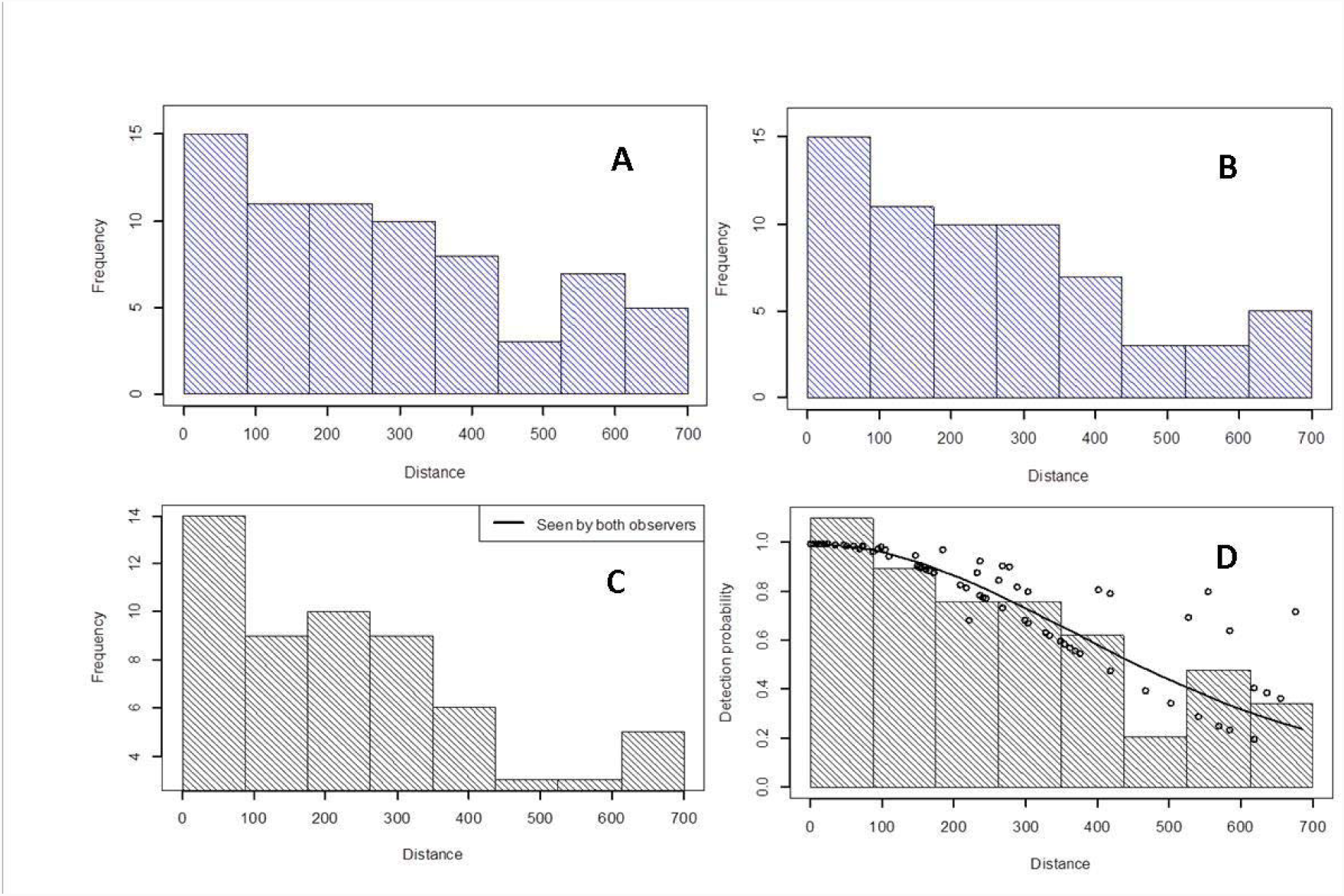
Scaled histograms of perpendicular distances and the predicted detection probability functions for fin whale groups. A) Front observers, B) rear observers, C) both observers and D) pooled detections with sightings fitted to a half-normal model where each observation prediction is shown as a circle. Distance is measured in meters from the line directly below the aircraft.

The MRDS analysis provided partially corrected estimates of fin whales of 465 (95% CI: 233-929) and 1,932 (1,204-3,100) in West and East Greenland, respectively (Table 5).

**Table 5.** Development of availability correction factors for minke, fin and humpback whales for different distance truncations for detections from 0 to 2 m depth. The cue rates were obtained from Heide-Jørgensen and Simon (2007). The beta distribution of the surface availability (see Table 1) was skewed to the right and restricted to avoid unrealistically low (i.e.<12%) surfacing times. A Kolomogorov-Smirnov test detected a significant difference for the time-in-view distributions of fin whales from East and West Greenland in 2015 (p=0.013) but no difference was found for humpback whales (p=0.872) or minke whales (p=0.604) for the same year.

| Distance truncation | Time-in-view, Mean, range, n | Surface availability, | Cue rate $\text{hr}^{-1}$, Mean, SD | Availability correction factor, Mean, cv |
| --- | --- | --- | --- | --- |
| Minke whale East and West Greenland 2015 |  |  |  |  |
| <450 m | 2.9, 0-14 s, 40 | 2.5, 4, 12, 23 | 46.3, 10.9 | 0.19, 0.27 |
| Minke whale West Greenland 2007 |  |  |  |  |
| 240 m | 5.6, 0-15 s, 8 | 2.5, 4, 12, 23 | 46.3, 10.9 | 0.21, 0.25 |
| Humpback whale 2015 |  |  |  |  |
| 50-1200 m | 8, 0-60 s, 124 | 2.5, 4, 22, 45 | 71, 5 | 0.43, 0.27 |
| Fin whale East Greenland 2015 |  |  |  |  |
| <700 m | 11.3, 0-53 s, 113 | 2.5, 4, 15, 25 | 50, 3.5 | 0.30, 0.10 |
| Fin whale West Greenland 2015 |  |  |  |  |
| <700 m | 3.4, 0-15 s, 20 | 2.5, 4, 15, 25 | 50, 3.5 | 0.21, 0.22 |
| Fin whale 2007 |  |  |  |  |
| 50-800 m | 6, 0-12 s, 9 | 2.5, 4, 15, 25 | 50, 3.5 | 0.28, 0.22 |
| Fin whale 2005 |  |  |  |  |
| <2500 m | 12, 0-63 s, 56 | 2.5, 4, 15, 25 | 50, 3.5 | 0.33, 0.43 |

The observed surface time for the fin whales tracked in West Greenland was 19.45% (Table 1) and the mean time-in-view of fin whale sightings was 11s and 4s in East and West Greenland, respectively (Table 5). Heide-Jørgensen and Simon (2007) observed that fin whales in West Greenland had a blow rate of 50 times per hour (cv=0.07) when excluding observation periods <30min. This corresponds to an average duration of surfacings per hour of 13.1s (3,600*0.20/50) and an average duration of dives of 58.9s (3,600-(1-0.20))/50). Using these values in the model by Laake et al. (1997) results in an availability for fin whales of 0.30 (cv=0,10) and 0.21 (cv=0.22) in East and West Greenland, respectively (Table 5). Applying this to the MRDS estimates give fully corrected abundance estimates of 6,440 (95% CI: 3,901-10,632) and 2,215 (95% CI:1,017-4,823) fin whales in East and West Greenland, respectively (Table 7).

Fin whale abundance estimates from 2005 and 2007 were not corrected for availability bias but applying the same availability bias as for the 2015-survey, corrected for the specific time-in-view data from 2005 and 2007, provided fully corrected abundance estimates of 9,800 (95% CI: 3,228-29,751) in 2005 (Heide-Jørgensen et al. 2008) and 15,957 (95% CI 4,531-56,202) in 2007 (Heide-Jørgensen et al. 2010b).

#### Humpback whale abundance

In West Greenland most humpback whales were detected in the southern strata, but densities were low compared to East Greenland where humpback whales were detected in large numbers all along the coast (Fig. 7). Still, the number of sightings in West Greenland was similar to that detected in the survey in 2007 (Table 3).

**Fig. 7.**
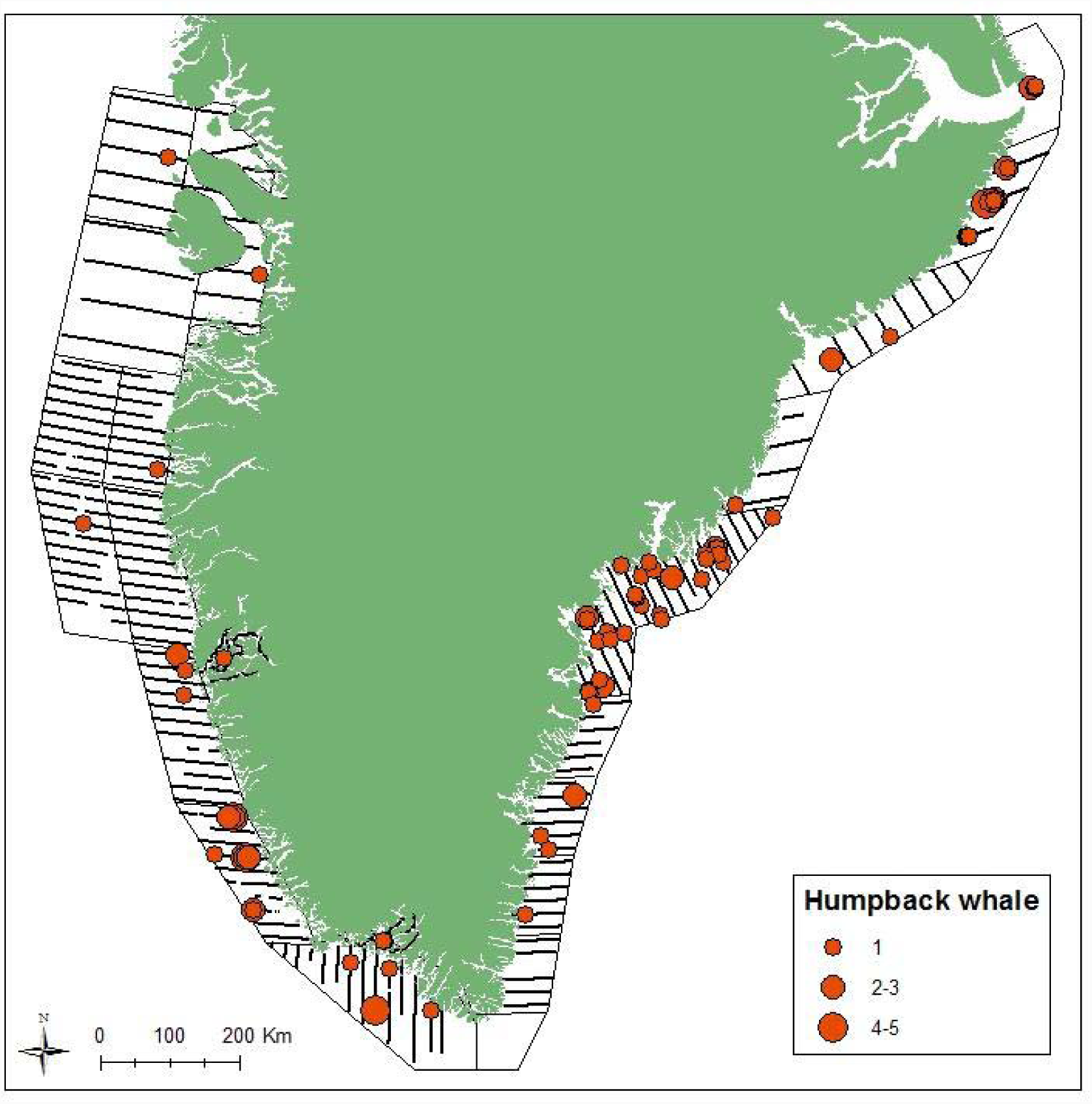
Survey effort in sea states <5 and sightings with group sizes of humpback whales in East and West Greenland.

The number of sightings of humpback whales in sea states <5 was almost three times higher in East Greenland compared to West Greenland (Table 2). The northernmost stratum in East Greenland (stratum E1, Figs 1 and 7) had 7 sightings of humpbacks but the coverage was restricted to one line where only half the line (23 km) was covered in good survey conditions. Because of the biased coverage of stratum E1 abundance was not estimated for this stratum.

The overall expected group size of humpback whales was 1.35 (cv=0.09) in East and 1.53 (cv=0.16) in West Greenland.

The average time-in-view for both observers for humpback whales was 8 s (n=124). Adjusting the average surface time (33.5%, cv=0.10, Heide-Jørgensen and Laidre 2015) for the time-in-view factor using the approach from Laake *et al.* (1997) gives an appropriate availability bias correction of 0.43 (cv=0.27) for both East and West Greenland in 2015 (Table 4).

The MRDS analysis was right truncated at 1200 m and left truncated at 50 m leaving 76 observations for abundance estimation. The explanatory variables available to be included in the MRDS models were, in addition to perpendicular distance to sightings; group size, sea state and observer. The final MRDS model included observer as a variable because the front observers had more sightings (Fig. 8). The MRDS analysis provided partially corrected estimates of humpback whales of 1816 (95% CI: 933-3536) and 427 (219-831) in East and West Greenland, respectively (Table 5). There was little perception bias in the combined East and West Greenland data(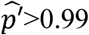, Table 4) and the MRDS analysis with separate East and West Greenland global group size estimates provided fully corrected estimates of humpback whales of 4,223 (1,845-9,666) and 993 (95% CI: 434-2,272) in East and West Greenland, respectively (Table 7).

**Fig. 8.**
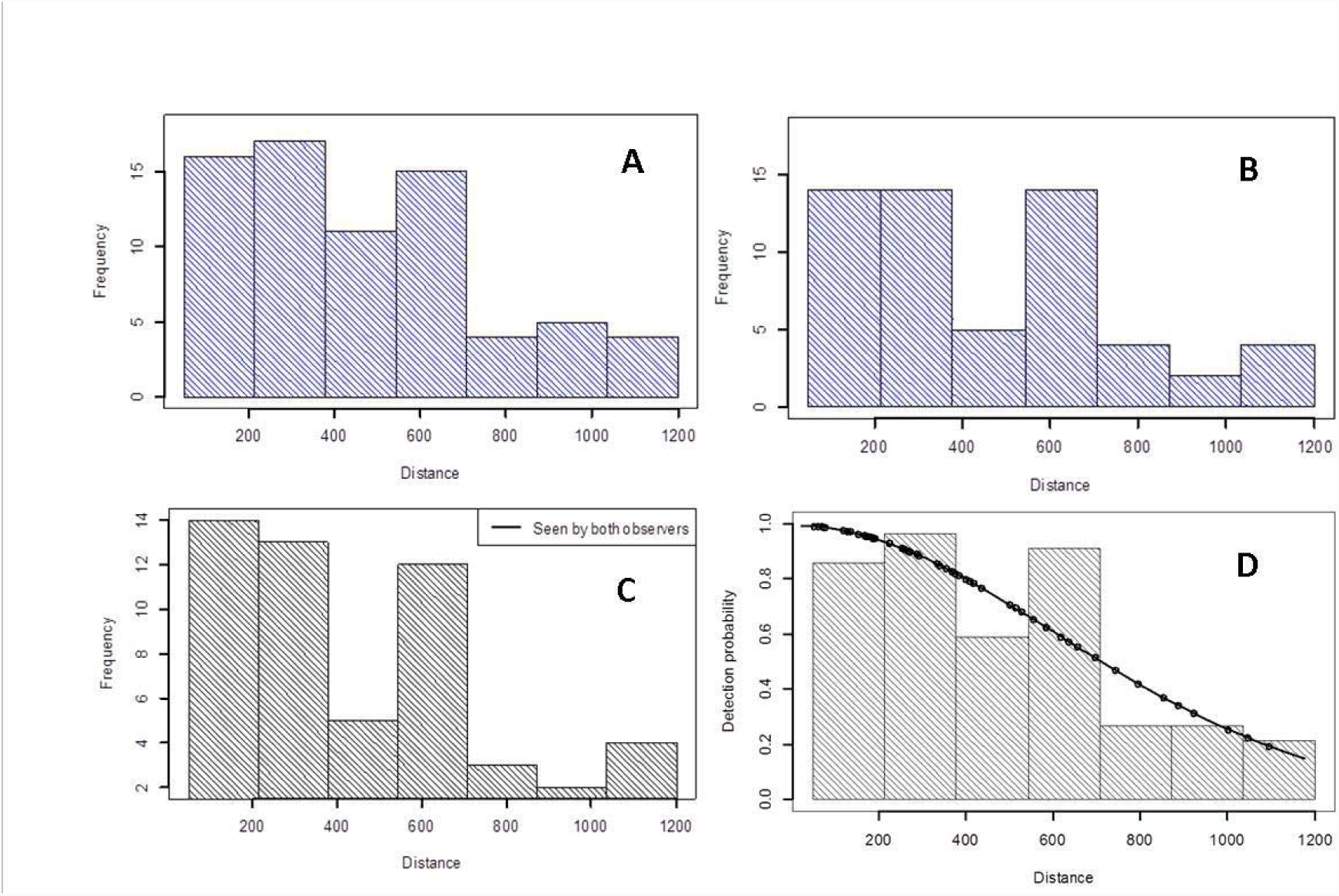
Scaled histograms of perpendicular distances and the predicted detection probability functions for humpback whale groups. A) Front observers, B) rear observers, C) both observers and D) pooled detections with sightings fitted to a half-normal model where each observation prediction is shown as a circle. Distance is measured in meters from the line directly below the aircraft. The data were left truncated at 50m due to low number of detections close to the trackline.

#### Harbour porpoise abundance

Except for two sightings in East Greenland harbour porpoises were only detected in West Greenland and there were more sightings in 2015 compared to 2007 (Fig. 9, Table 3). A right truncation at 250 m left 48 observations in sea states<3 for abundance estimations. A half-normal key with no covariates was chosen for the MRDS model (Fig. 10) that provided at-surface abundance estimates of 15,831 harbour porpoises (95% CI: 8,514-31,202, Table 6) in West Greenland with a perception bias of 0.82 (cv=0.21, Table 4). The overall expected group size of harbour porpoises was 2.00 (cv=0.71) in East and 1.79 (cv=0.09) in West Greenland.

**Fig. 9.**
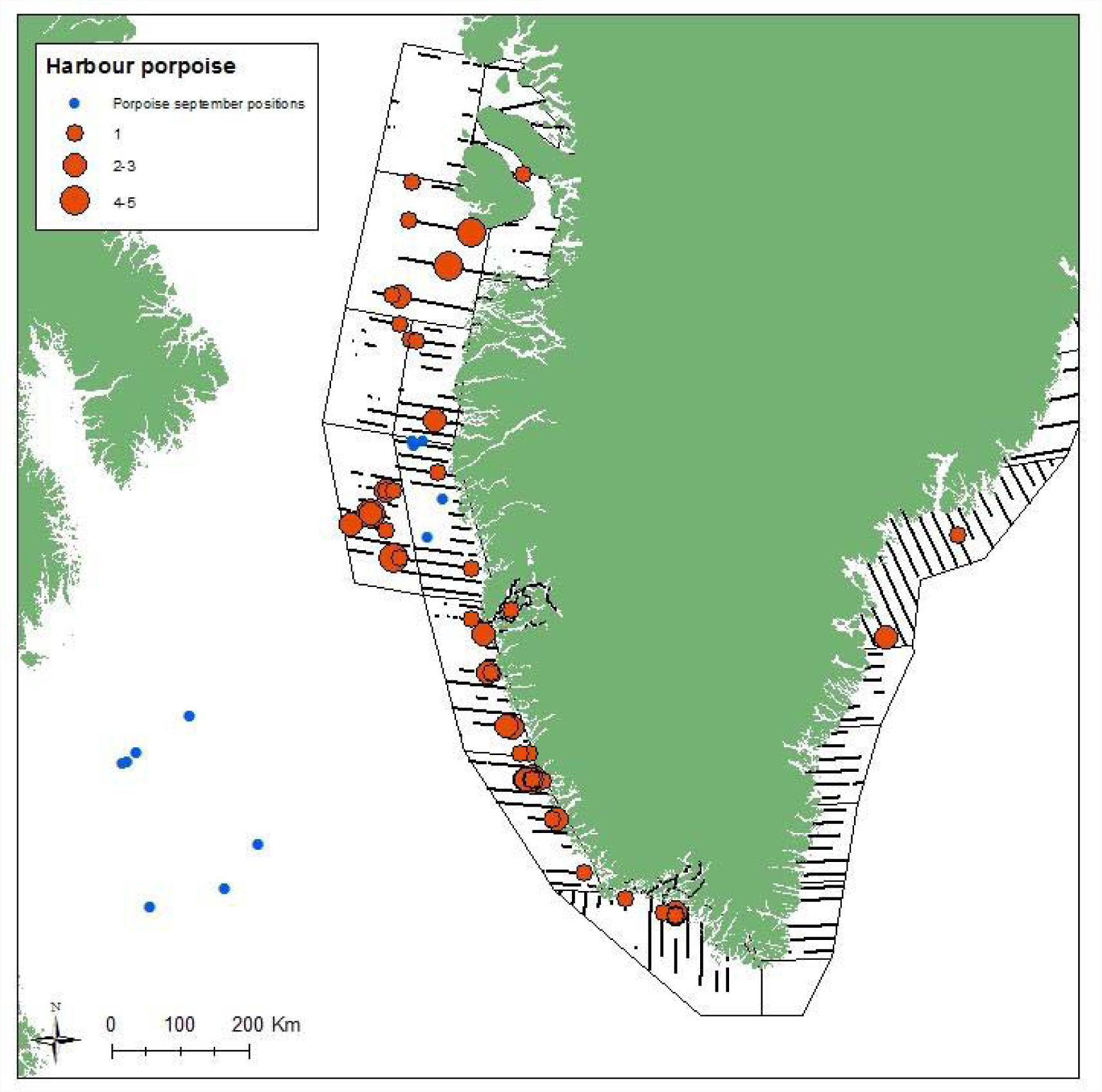
Survey effort in sea states <3 and sightings with group sizes of harbour porpoises in East and West Greenland. Blue dots indicate satellite positions of harbour porpoises tagged inside the survey area and tracked in September 2015.

**Fig. 10.**
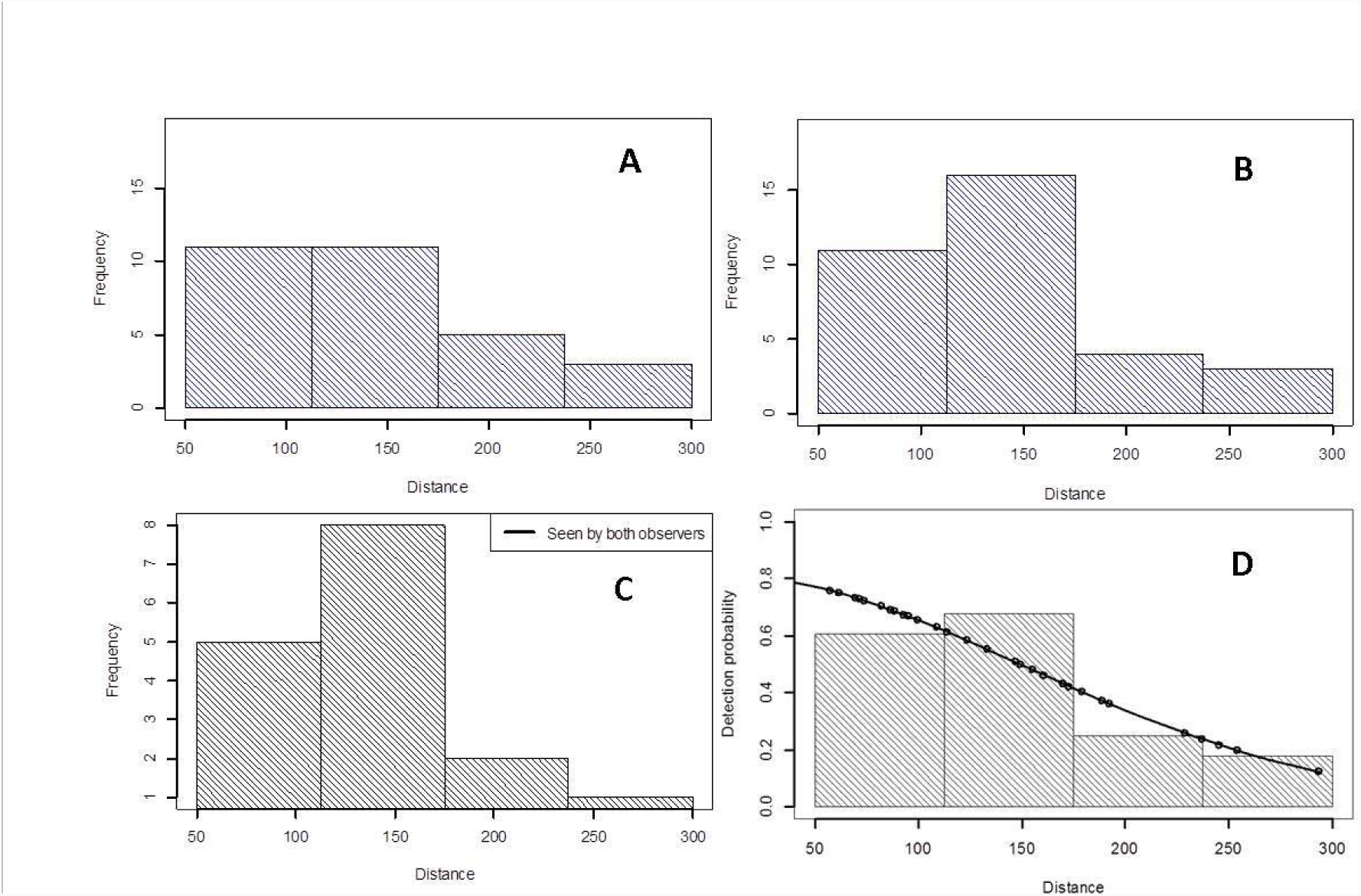
Scaled histograms of perpendicular distances and the predicted detection probability functions for harbour porpoise groups. A) Front observers, B) rear observers, C) both observers and D) pooled detections with sightings fitted to a half-normal model where each observation prediction is shown as a circle. Distance is measured in meters from the line directly below the aircraft.

The average time-in-view for harbour porpoises was 0.8 s and it was considered too low for being relevant for adjusting the availability bias. Based on satellite tracking of 9 harbour porpoises in West Greenland an availability correction factor was developed (Table 1) and applying this to the at-surface abundance gave a fully corrected estimate of 83,321 harbour porpoises (95% CI: 43,377-160,047, Table 7) in West Greenland and 1,642 (95% CI: 319-8,464) in East Greenland. Applying this new developed correction factor to the harbour porpoise at-surface estimate from 2007 gave a fully corrected estimate of 54,284 harbour porpoises (95% CI: 27,627-106,664).

#### Pilot whale abundance

Except for four sightings in East Greenland pilot whales were only detected in West Greenland (Fig. 11). More pilot whales groups were detected in 2015 than in 2007 (Table 3). A right truncation at 700 m left 32 observations in sea states<5 for abundance estimations. The expected group size was 8.5 whales (cv=0.10). A half-normal key with no covariates was chosen for the MRDS model (Fig. 12) that provided at-surface abundance estimates of 4,797 pilot whales (95% CI: 1,793-12,832) in West Greenland and 135 (16-1,142) in East Greenland with a perception bias of 0.98 (cv=0.029, Table 4 and 6). Based on satellite tracking of 2 pilot whales in the Faroe Isles in 2001 (Heide-Jørgensen et al. 2002) an availability correction factor of 0.40 (cv=0.15, Hansen and Heide-Jørgensen 2013) was developed (Table 1). Applying this factor to the at-surface abundance gave a fully corrected estimate of 9,190 pilot whales (95% CI: 3,635-23,234) in West Greenland and 258 (95% CI: 50-1,354) in East Greenland (Table 7). Pilot whales are however detectable ahead of the plane and applying an instantaneous availability correction factor leads to a positive bias. The average time-in-view for pilot whales was 6.4 s (cv=0.76) but data on the number of surfacings/dives are missing and adjustment of the availability bias for time-in-view is not possible. For a comparison with the survey in 2007 it is still useful to apply the same availability correction factor to the survey in 2015.

**Fig. 11.**
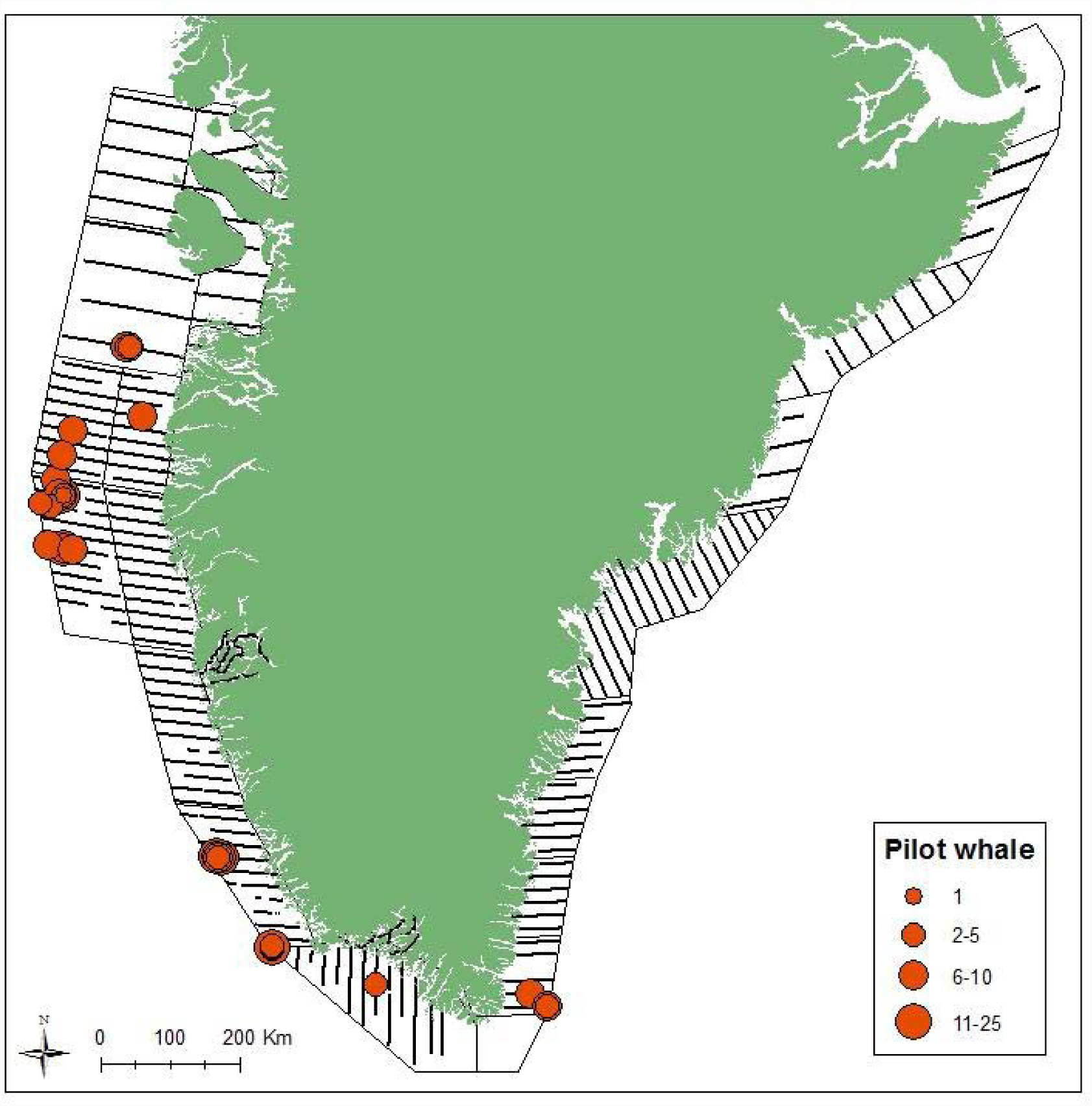
Survey effort in sea states <5 and sightings with group sizes of pilot whales in East and West Greenland.

**Fig. 12.**
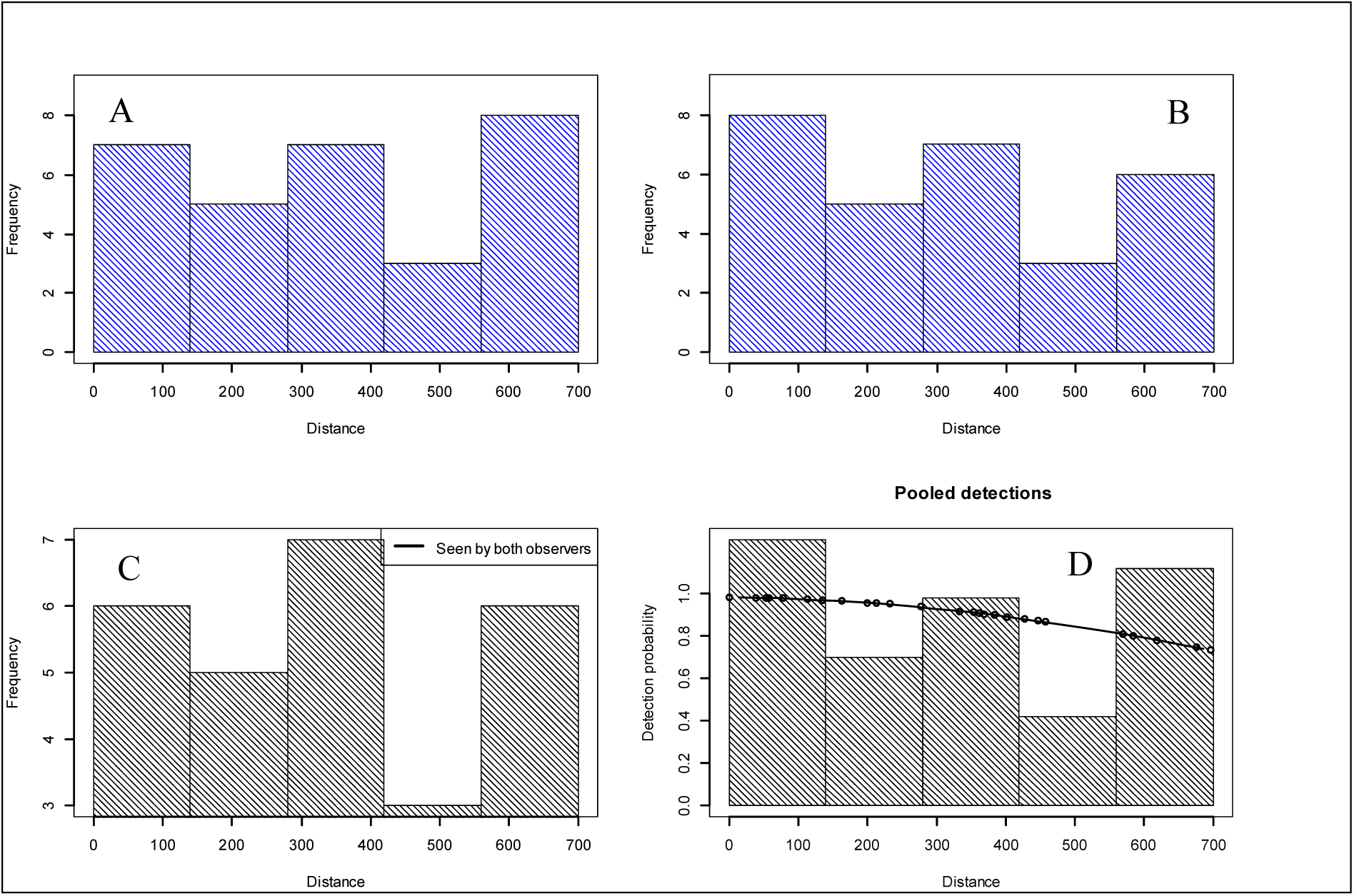
Scaled histograms of perpendicular distances and the predicted detection probability functions for pilot whale groups. A) Front observers, B) rear observers, C) both observers and D) pooled detections with sightings fitted to a half-normal model where each observation prediction is shown as a circle. Distance is measured in meters from the line directly below the aircraft.

#### White-beaked dolphin abundance

White-beaked dolphins were widespread in both East and Southwest Greenland (Fig. 13) but the number of sightings in West Greenland in 2015 was only half of the sightings in 2007 (Table 3).

**Fig. 13.**
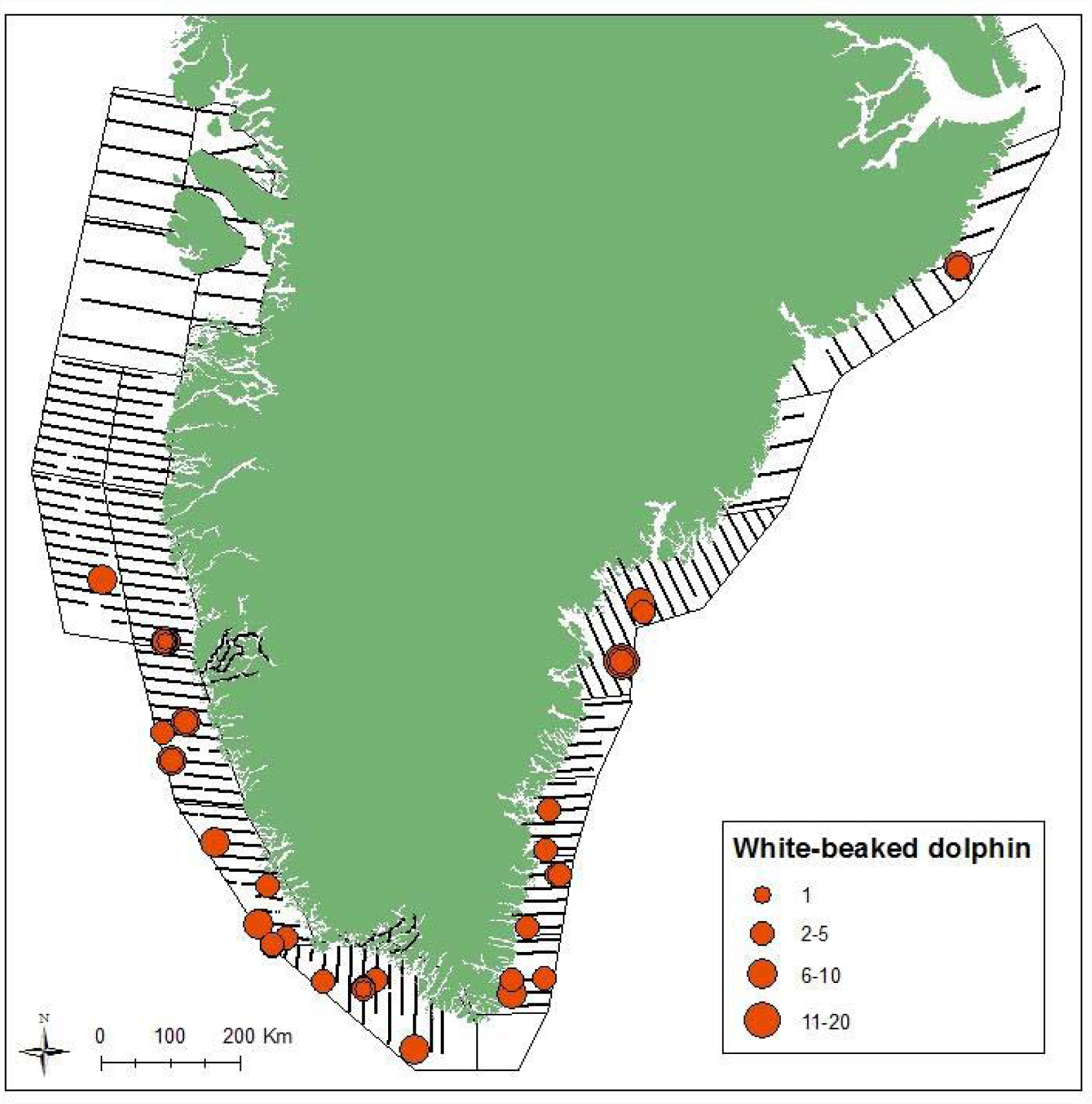
Survey effort in sea states <5 and sightings with group sizes of white-beaked dolphins in East and West Greenland.

The expected group size was 4.2 (cv=0.19) in West Greenland and 4.5 (0.19) in East Greenland. A half-normal key with sea state as a variable in the DS component was chosen for the MRDS model (Fig. 14) that provided at-surface abundance estimates of 2,747 white-beaked dolphins (95% CI: 1,257-6,002) in West Greenland and 2,140 (95% CI: 825-5,547) in East Greenland with a joint perception bias of 0.99 (cv=0.01, Table 4).

**Fig. 14.**
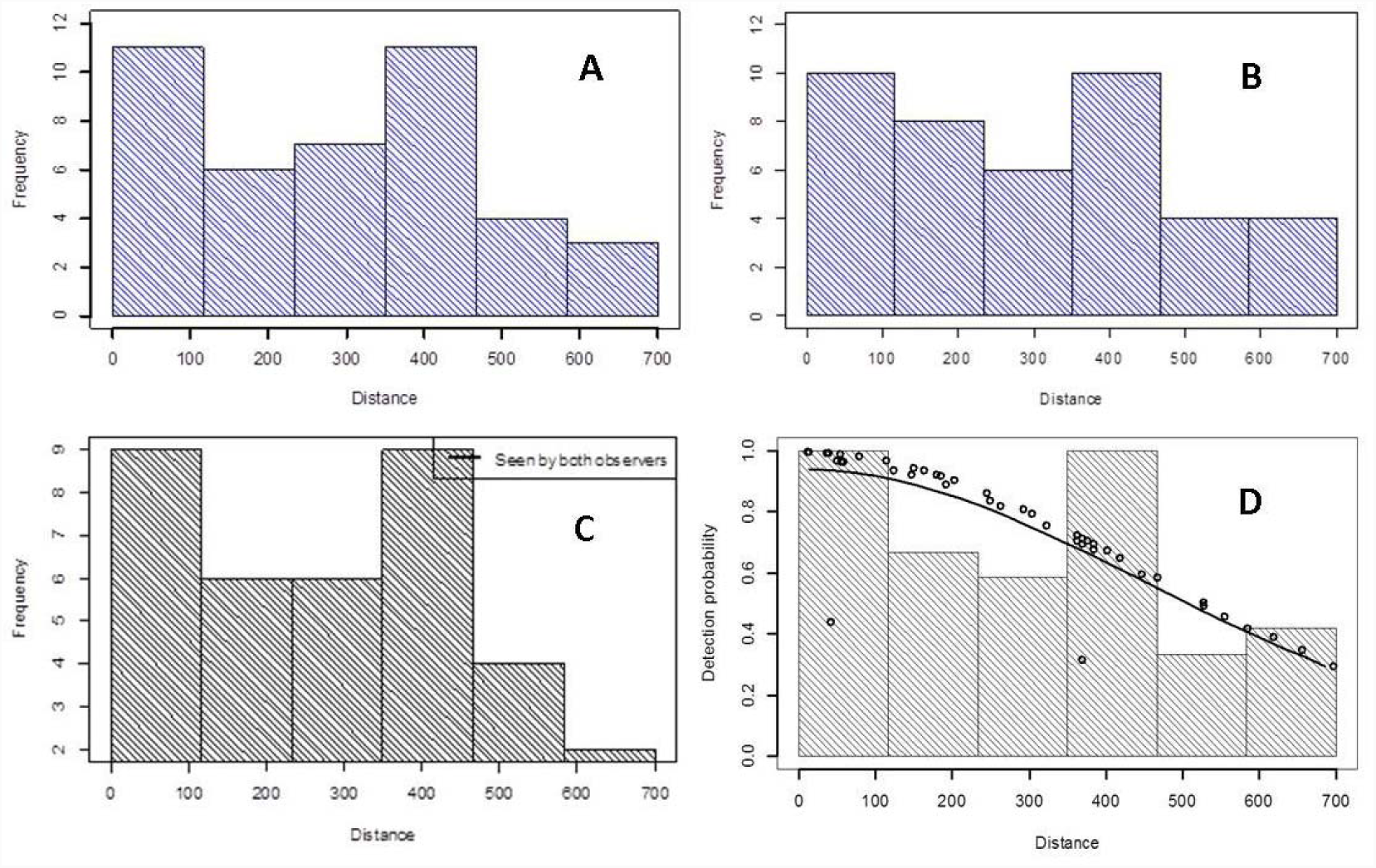
Scaled histograms of perpendicular distances and the predicted detection probability functions for white-beaked dolphin groups. A) Front observers, B) rear observers, C) both observers and D) pooled detections with sightings fitted to a half-normal model where each observation prediction is shown as a circle. Distance is measured in meters from the line directly below the aircraft.

Hansen and Heide-Jørgensen (2013) used data from a single white-beaked dolphin from Iceland to develop an availability correction factor (Table 1) and applying this to the at-surface abundance gave a fully corrected estimate of 15,261 white-beaked dolphins (95% CI: 7,048-33,046) in West Greenland and 11,889 (4,710-30,008 in East Greenland (Table 8). As for pilot whales the availability correction factor is not adjusted for time-in-view but for a comparison with the survey in 2007 it is useful to apply the same availability correction factor.

#### Detection of other species

Bottlenose whales, *Hyperoodon ampullatus*, (group size 1-4) were seen in West Greenland and sperm whales, *Physeter catodon,* (1-2) and killer whales, *Orcinus orca*, (1-5) were seen both in East and West Greenland (Figs 15). One sei whale, *Balaenoptera borealis*, was seen in West Greenland and one blue whale, *Balaenoptera musculus*, and one group of five narwhals, *Monodon monoceros*, were seen in East Greenland (Fig. 15).

**Fig. 15.**
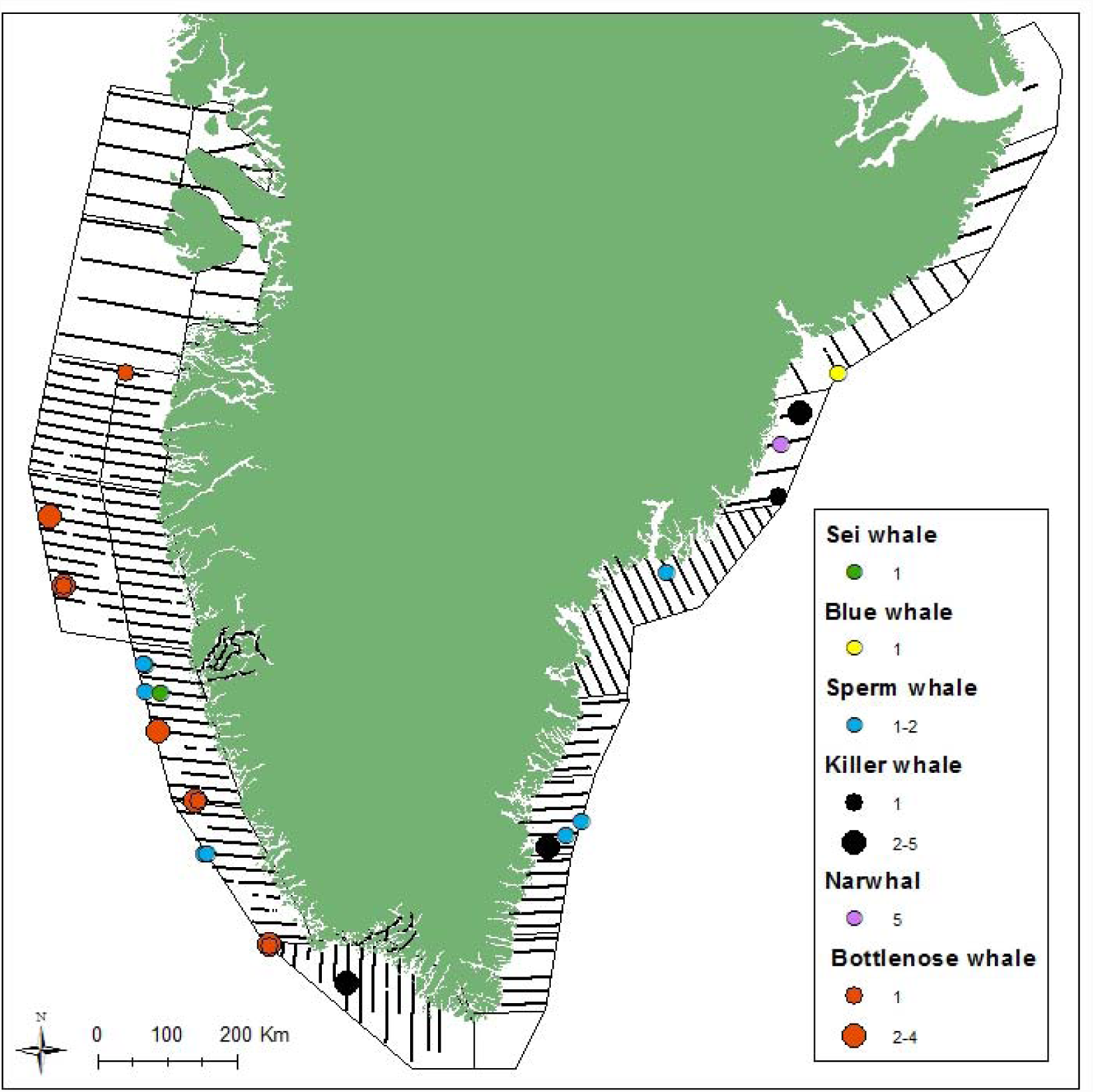
Survey effort in sea states <5 and sightings with group sizes of narwhals, sei, blue, sperm, bottlenose and killer whales in East and West Greenland.

## DISCUSSION

MRDS estimates that correct for perception bias are considered the most accurate estimates for all species as they take into account the perception bias and the heterogeneity that affects the observers’ detections of the whales. Due to the low number of sightings of minke whale in both 2007 and 2015 precluded the use of MRDS methods. The number of sightings of minke whales has generally been low in all previous surveys of the area (Heide-Jørgensen *et al* 2010a). Realistic abundance estimates are dependent on correction for availability as minke whales spend only 16% of the time in the top 2 m of the water column where they would be visible to aerial observers (Table 1).

We therefore estimated abundance of minke whales using a strip census with an assumed width of 300 m for West Greenland, an approach that is consistent with the 2007 estimate for the same area (Heide-Jørgensen *et al.* 2010a). For East Greenland, we estimated abundance of minke whales using two approaches: 1. An MRDS analysis using the combined detection function from East and West Greenland, and 2. a strip census using an assumed strip width of 450 m. The wider strip width for East Greenland was chosen based on the observed detections for that area (Fig. 4). Both strip census estimates likely suffer from negative bias due to the assumption of constant detection across the strip width, although this assumption is supported by the observed detection functions. The strip transect estimate for East Greenland is also likely negatively biased because no correction for perception bias could be applied. Given these considerations, we prefer and present the estimates from the strip transect analysis for West Greenland, and the combined East/West MRDS analysis for East Greenland.

Fin whales and white-beaked dolphins have availability correction factors developed from instrumentations of only one or two individuals and these correction factors should be considered provisional, and until more samples become available it must be emphasized that the uncorrected MRDS estimates are negatively biased.

Satellite-linked time-depth-recorders may have a slight drift in the pressure transducers (Heide-Jørgensen and Laidre 2015) but a more important factor is the position of the instruments on the whales as that determines when the surface is reached and the salt water switch is activated. Since the transmitters were remotely deployed on minke and fin whales, the position of the tag on the whale cannot be controlled with precision and may even change while the transmitter is migrating out through the skin. To reduce the possible influence of tag position on the data collection it is important to evaluate the surface availability on larger sample sizes and the samples of five minke whales and two fin whales need to be increased.

Fin whales and white-beaked dolphins have availability correction factors developed from instrumentations of only one or two individuals and these correction factors should be considered provisional, and until more samples become available it must be emphasized that the uncorrected MRDS estimates are negatively biased.

The development of an adjustment of the availability correction for the time-in-view was based on cue-rates for the three baleen whales with fully corrected abundance estimates. For these species a cue is defined as any part of the body of the whale appearing at the water surface. There will however often be several cue’s involved in a surfacing event (the period between ascent above 2 m and before next descend below 2 m). It is therefore likely that the impact of the time-in-view correction is overestimated and the abundance estimates negatively biased. The magnitude of the bias is difficult to assess but it could be as large as 1-2 percentage points of the availability correction factor. A more accurate correction factor could be developed if data on the average duration of surfacings and dives were available.

The fully corrected abundance estimates of fin whales between 2005 and 2015 from West Greenland fluctuate more widely than the other species and this is mainly due to a major difference in the group size estimates that reached its maximum in 2007 with a mean observed group size of 2.5 (range 1-25) compared to 1.7 (range 1-13) in 2005 and 1.5 (range 1-7) in 2015. The large groups observed in previous years in West Greenland were not detected in West or East Greenland (mean=1.6, range 1-6) in 2015, and the changes are most likely due to ecological changes that has influenced the distribution of fin whales in the North Atlantic.

All of the three large whale species (minke, fin and humpback whale) show a remarkable decline in West Greenland since the last abundance estimates were obtained in 2007 (Fig. 16). The minke whale has decline from 10,000 to 5,000, the fin whale from 4,400 to 1500 and the humpback whale from 2,704 to 1,000. The same survey design and some of the same observers were used in the two surveys, and survey conditions were in both surveys kept to sea states below 3 for minke whales and below 5 for fin and humpback whales. Identical survey techniques were deployed in the two surveys and the decline cannot be attributed to a lower detection probability due to new observers in the 2015-survey because detections of large whales in East Greenland (that was covered first and could be considered as training) were still unexpectedly high. In addition, detection of harbour porpoises, which is the smallest and the most cryptic of all the whale species, were similar or higher in the 2015 West Greenland survey compared to the survey in 2007. Thus the observed changes in abundance of the large cetaceans and white-beaked dolphins cannot be considered an artefact of survey methods but must reflect a real decline in abundance in West Greenland.

**Fig. 16.**
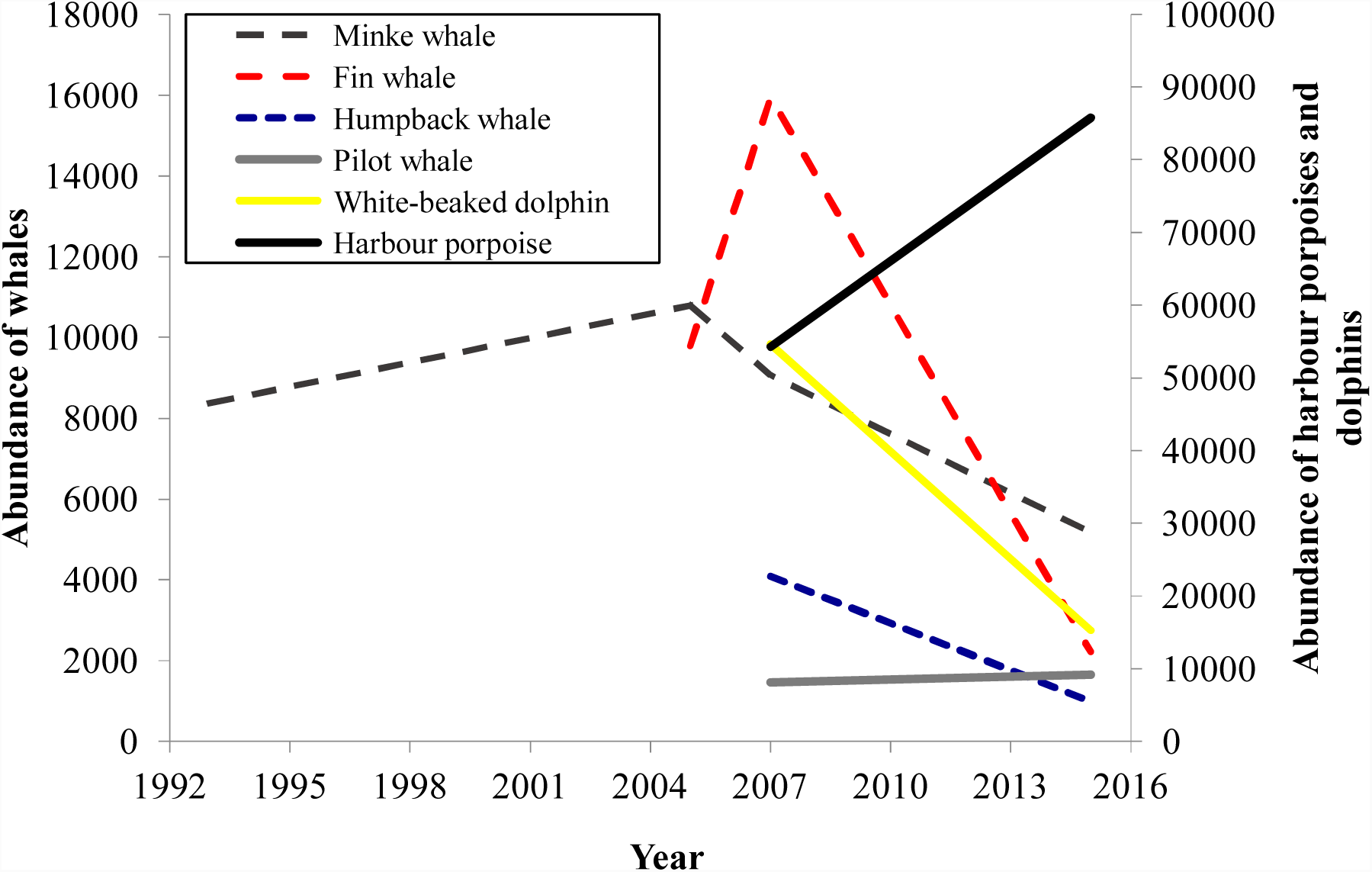
Trends in abundance of six cetacean species in West Greenland where compatible abundance estimates are available. The harbour porpoise and white-beaked dolphin abundances are shown on the right y-axis. Data from 2007 and earlier from Hansen and Heide-Jørgensen (2013), Heide-Jørgensen and Laidre (2015), Heide-Jørgensen et al. (2010a and 2010b).

The at-surface abundance estimate of harbour porpoises in West Greenland is slightly larger but not significantly different from the estimate from the 2007-survey (10,021, cv=0.31). Satellite tracking of harbour porpoises in 2012-13 has shown that during August-September they only spend about 73% of their time inside the survey strata in West Greenland (GINR unpublished data). In September 2015, the positions from two harbour porpoises tagged in West Greenland were available and one of the porpoises had left the Greenlandic coastal areas at the time of the survey suggesting that only 50% of the population was within the surveyed area (GINR and Nielsen et al. in press). Annual variations in the proportions of harbour porpoises that are within the surveyed area are likely to explain the variability in the abundance estimates and further indicate that the ecological changes that have affected the presence of large whales in West Greenland have not affected the abundance of harbour porpoises.

The estimate of pilot whales in West Greenland is similar to the estimate from 2007. The survey was designed to cover the distribution of the baleen whales in West Greenland and hence did not cover the most westerly part of the pilot whale distribution. The deeper parts of the strata that pilot whales prefer was surveyed more intensively in 2015 than 2007 (Hansen and Heide-Jørgensen 2013). Pilot whales are, like harbour porpoises, also present further west in the deep waters in Baffin Bay and annual variations of the influx to the surveyed area may explain the variations in the abundance estimates.

Finally, the white-beaked dolphins are also showing a decline in abundance in West Greenland that may be related to the same ecological changes that apparently are making East Greenland a more favourable habitat for baleen whales. In 2007 more dolphins were detected in the northern strata (Hansen and Heide-Jørgensen 2013) and although species identification was uncertain it was most likely white-beaked dolphins that in 2007 were more widespread along the West Greenland coast.

The east coast of Greenland is periodically blocked by dense masses of multi-annual drifting pack ice that is produced in the Arctic Ocean and is transported by the current through Fram Strait south along the coast of Greenland. With the exception of Scoresby (1823) the east coast of Greenland was rarely or never visited by whalers that were pursuing bowhead whales. The main reason being the severe border of drifting pack-ice along the coast that prevented the entry of whalers to the coast. In the 19th century the east coast of Greenland was only inhabited in the Tasiilaq area by a few hundred Inuit hunters. Based on examination on expedition reports, local informants and official accounts from the Royal Greenland Trade Department Winge (1902) concluded that are no confirmed information about humpback whales and fin whales in East Greenland. Holm and Petersen (1921) and Jensen (1928) mentioned that humpback and fin whales visit coastal areas of Tasiilaq infrequently and only in years where no coastal ice is present in summer. Born (1983) mention that two humpback whales were seen in Scoresby Sound in August 1980.

There are no previous abundance estimates from the East Greenland coastal area and the estimates of all three large cetaceans from the relatively small area (half the size of the area in West Greenland) seem surprisingly high considering that very few observations of baleen whales have been recorded in coastal areas of East Greenland for the past 150 years. Offshore areas along the coast of East Greenland have been surveyed in the past and large numbers of especially fin whales have been detected between East Greenland and Iceland (Vikingsson et al. 2009). It is possible that whales, that previously were feeding further offshore, now have a more coastal distribution in summer, perhaps facilitated by the reduction of summer sea ice in East Greenland in the past decade (Kern et al. 2010).

The mackerel (*Scomber scombrus*) fishery in East Greenland has increased dramatically since 2011 when the first catches of <1 ktons were recorded up to 2014 when more than 78 ktons were caught (Jansen et al. 2016). The rapid expansion of mackerel to the East Greenland shelf areas is related to an increased summer SST with temperatures above 6°C. Even though mackerel is not known to be a primary prey item for baleen whales it seems likely that the dramatic increase in a pelagic prey resource may be driven by some of the same factors that are also driving baleen whale and dolphin distribution (e.g. krill, copepods. Myctophidae and herring, *Clupea harengus*). Large-scale ecological changes on both sides of Greenland are most likely driving the observed shifts in abundance of whales in the two areas.

## ACKNOWLEDGEMENTS

This study was funded by the Greenland Institute of Natural Resources, the Greenland Government (Department for Fish and Wildlife) and the North Atlantic Marine Mammal Commission. We wish to thank Norlandair for providing aircrafts and skilled pilots and for excellent collaboration during all phases of the surveys.

